# Precise Extraction of Neural Motifs Reveals Multiscale, Parallel Encoding Schemes in Auditory Cortex

**DOI:** 10.1101/2025.08.26.672281

**Authors:** Liang Xiang, Mingxuan Wang, Patrick O. Kanold, Adam Charles

## Abstract

Neural population activity is often stereotyped into recurring activity patterns, i.e., neural motifs, which can be seen as the fundamental building blocks in sensory processing and cognition. In this work we study the codes carried by such neural motifs in primary auditory cortex A1, and analyze how they build on and complement traditional views of single-unit coding. In particular, we study, using two-photon calcium imaging (CI), how activity in A1 differentially represents both sensory stimuli and task and behavioral variables in each of the parallel single neuron and population motif scales. While CI enables the study of neural motifs by capturing the activity of large neural populations, identifying motifs in CI is hampered by the temporal imprecision of current motif-detection algorithms when applied to CI data. We thus developed a new algorithm for motif detection, which enabled us to identify widespread stimulus-encoding motifs as well as a small number of motifs jointly encoding stimulus and choice in Layer 2/3 of A1. These motifs consist of small groups of neurons that are neither clustered nor regularly ordered in space. Interestingly, active neurons within task-encoding motifs exhibit mixed encoding properties inconsistent with the motifs they participate in. Together, these results demonstrate how single unit activity and neural motifs in A1 L2/3 provide different levels of coding granularity containing different information in parallel within the greater neural population. Generally, our results indicate that downstream populations, by selecting which scale of a population drives them, can be selective in the information collected for later cognitive processing.

## 1 Introduction

Navigating and interacting with the external world requires that organisms process massive amounts of sensory information to guide behaviors. Understanding how sensory and behavioral information are encoded and transmitted by neural populations is one of the most fundamental problems in system neuroscience. Traditionally, neural coding has been considered in terms of single-cell coding, e.g., via tuning curves^(1;2;3;4)^ or receptive fields^(5)^, where a single neuron encodes and conveys a given piece of information. Alternatively, it has been hypothesized that such encoding and transmission of information in cortical circuits may be accomplished by stereotypical and recurring neural firing patterns, i.e, neural motifs, which have also been called synfire chains, sequences, packets, ensembles, etc.^(6;7;8;9;10)^. These motifs, unlike single cell activity, capture the dynamic flow of neural activity through networks and have been considered as vital neural computational building blocks in sensory processing and cognition. Understanding how information encoded in motifs complements single-neuron coding would define how population-level dynamics adds new layers of accessible, independent information to downstream areas.

Neural motifs have been reported in many brain regions across multiple species over a wide range of settings, including sensory processing^(8;11;12)^, motor production^(13;14)^, memory replay^(15;16;17)^, and decision making^(18)^. In early sensory cortices, the discovery of spontaneous and stimulus-evoked neural motifs has exposed the complex processing of sensory information by neural populations^(8;9)^. However, more recent experiments have shown that neural activity in sensory cortices is not only representative of the sensory stimulus but is also modulated by behaviors and contexts^(19;20;21;22;23;24;25)^. In fact, large-scale neural recordings have revealed distributed coding of task and behavioral variables, e.g., stimulus, motion, decision and engagement, across the entire brain, including primary sensory areas^(26;27)^. It is unknown whether these non-traditional task-encoding signals are fundamentally ingrained, i.e., they are a part of stereotyped neural motifs.

Recent use of in vivo calcium imaging enabled the large-scale discovery of motifs^(28)^. Studies associating neural population activities with decisions, indirectly support the presence of choice-encoding motifs in the auditory cortex^(29;30;31)^. Specifically, Granger causality analysis showed that sound information used to inform decisions is transiently carried by individual neurons and transmitted sequentially within a small sub-network of neurons in the L2/3 of primary auditory cortex (A1)^(29;30)^. However, the presence and properties of non-auditory encoding motifs in the auditory cortex and the cellular composition of such motifs at single neuron level are unknown. We here investigated if both stimulus-encoding and choice-encoding neural motifs exist in A1 L2/3.

To identify motifs from large neuron population, one class of unsupervised methods based on Convolutional Non-negative Matrix Factorization (CNMF) with regularization ^(32;33;34;35;36)^ has been applied to spiking data, e.g., in identifying motifs of precise spike times in zebrafinch recordings ^(37)^. While these methods have subsequently been applied to widefield calcium imaging data^(28)^, current CNMF models using multiplicative updating rules (MUR) suffer low temporal precision in identifying motif onset times due to the temporal blurring in calcium imaging. Moreover, MUR fitting procedures are susceptible to low convergence rates, poor local minimum, and nonparsimonious solutions^(38;39)^, with alternative updating rules developed in the non-convolutional NMF domain to mitigate these effects^(40;39;41;42;43)^. Given that calcium imaging approaches can simultaneously record from larger neuron populations (hundreds to thousands of neurons) with precise cell localization, developing CNMF algorithms to appropriately process calcium imaging data would extend current motif identification to a finer temporal and spatial scale.

To more robustly study motifs in calcium imaging datasets, we first devised a new motif learning algorithm based on Split Bregman Iterations^(44;45)^ for sequence NMF (seqNMF), a CNMF optimization with regularization terms designed for neural data^(37)^. We then demonstrated on simulated neural data that our algorithm can identify motifs with higher temporal precision and is robust to various noise types. To identify the existence of auditory and non-auditory motifs, we then applied our motif detection algorithm to calcium imaging experiments of mice performing auditory tasks with different behavioral contingencies. We used a published two-photon calcium imaging dataset of mice performing an auditory go/no-go tone discrimination task^(29)^ and also imaged L2/3 neurons in mice performing a two-alternative forced-choice (2AFC) tone discrimination task.

In both datasets, we observed widespread presence of stimulus-encoding motifs and occasional occurrences of joint(stimulus and choice)-encoding motifs. When compared to the single neuron encoding properties, we found that individual neurons within a motif have mixed encoding properties that do not always align with the motif’s encoding properties, indicating that the encoding property of a single neuron does not necessarily represent how it is used at the population level. Despite the large number of auditory-encoding or movement-encoding single neurons, fewer than 10% are engaged in a single motif regardless of encoding property, indicating a sparse population code. Motifs can overlap, in that neurons can be shared between different motifs, especially between motifs with similar encoding properties. Together, these results reveal a parallel motif encoding scheme on a larger scale, which is fundamentally different from single-neuron encoding scheme in auditory cortex. The parallel streams open opportunities for downstream areas to access different pieces of information from the same neural population by accessing different levels of activity.

## 2 Results

### Stimulus and choice information are represented differently by neural population activities in A1

To test whether neuronal population activity formed motifs that include both sensory and nonsensory information, we studied population activity in the primary auditory cortex (A1) of mice, while animals performed tone discrimination tasks. As a preliminary analysis, we first tested whether stimulus and task-related information were represented differently by the neural population activity in different trials. We thus analyzed a previously collected two-photon calcium imaging dataset^(29)^ of neurons in mouse auditory cortex area Layer 2/3 in which mice were trained to discriminate different pure tones in a go/no-go paradigm^(29)^. Specifically, tones of four different frequencies (7kHz, 9.9kHz, 14kHz, 19.8kHz) were randomly played across trials. Mice were trained to lick a waterspout following a low-frequency target tone (7 or 9.9kHz) and to avoid licking in response to a high-frequency non-target tone (14 or 19.8kHz). Based on the frequency of the sound and behavioral response (correct or error), the trials were grouped by the 8 stimulus-choice combinations (Fig. 1A). If there exist stereotypical motifs encoding stimulus and behavior information, the neural population activity in the same trial types should be more similar than in different trial types. In 31 out of 34 sessions, we found that the cosine similarity scores (See STAR Methods) within similar stimulus trials were significantly greater than the similarity scores between different stimuli trials (Fig. 1B, Fig. S1, *χ*^2^ = 1.999 ∗ 10^3^, *p* < 1 ∗ 10^−8^, one-sided Wilcoxon rank sum test and Fisher’s combined probability test). Among the trials in which the same stimulus was played, the similarities within same choice trials were significantly greater than the similarities between different choices trials in 26 out of 34 sessions (Fig. 1B, Fig. S1, *χ*^2^ = 1.029 ∗ 10^3^, *p* < 1 ∗10^−8^, one-sided Wilcoxon rank sum test and Fisher’s combined probability test). The similar neural activity patterns within the same trial types demonstrate the joint encoding of stimulus and behavior information in A1 and suggest that motifs might exist, offering the potential for complementary encoding to known single-cell encoding properties.

**Figure 1.**
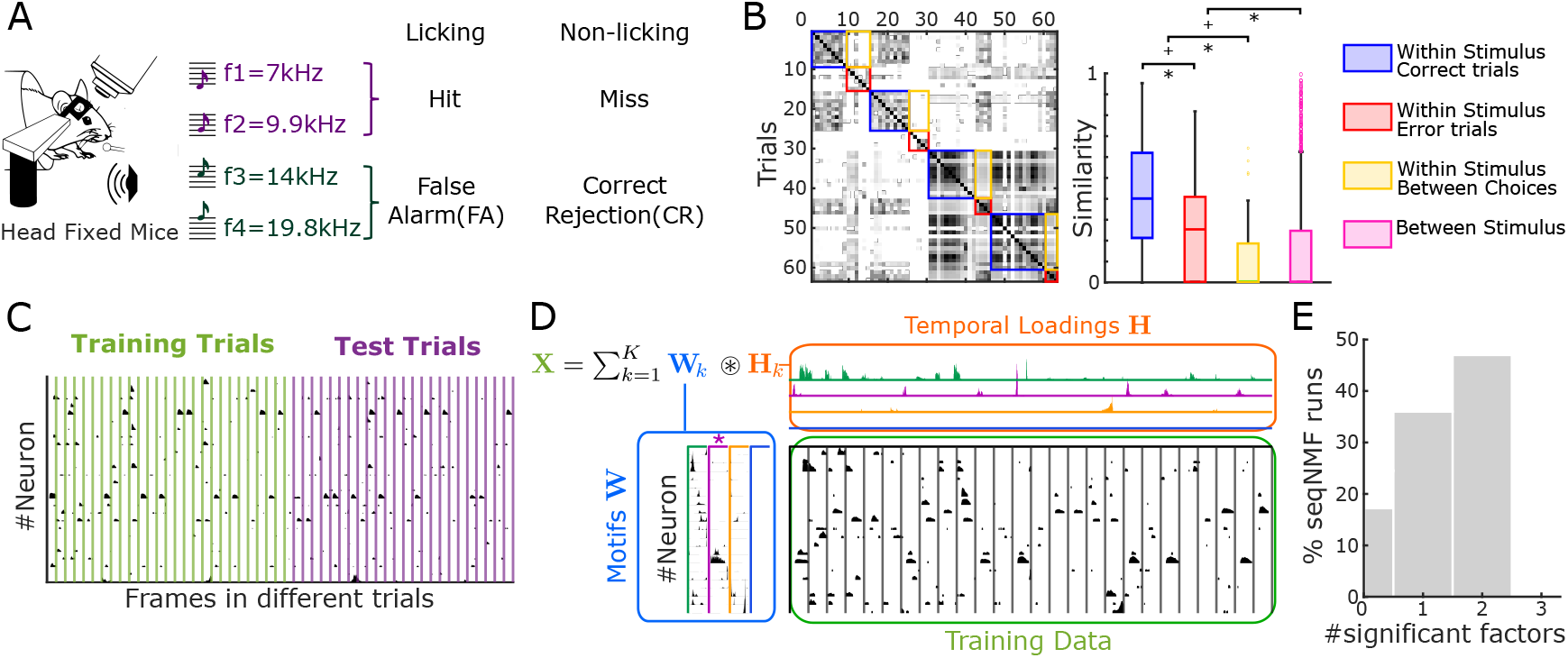
Finding motifs in A1 with Convolutional Non-negative Matrix Factorization (CNMF) (A) Experiment setup for mice performing a sound discrimination task. Mice were trained to lick the water spout in response to two low-frequency tones (7kHz and 9.9kHz) and avoid licking in response to two high-frequency tones (14kHz and 19.8kHz). The trial outcomes were categorized into hit, miss, false alarm(FA) and correct rejection(CR). (B) Similarity between neural activities in different trials within one example session, measured by the normalized dot products. Similarities were grouped by ‘within same stimulus, correct trials’, ‘within same stimulus, error trials’, ‘within same stimulus, between choices’ and ‘between stimulus’. Right: Box plot of the similarities in each group. (C) Trials were randomly split into training and test sets with balanced proportions of trials in the same frequency and choice groups (D) Schematic of SeqNMF. The input data matrix **X** (black) is decomposed into the sum of convolution of *K* different motifs (factors) **W** and their temporal occurrences **H**. Different motifs are marked with different colors (E) The number of significant motifs detected by SeqNMF on the same data was inconsistent, indicating SeqNMF factorization is sensitive to initialization.

### Current unsupervised motif discovery in calcium imaging lacks timing specificity

To precisely identify the potential motifs, we applied a recent motif detection algorithm designed for neural data, Sequence NMF (SeqNMF)^(37)^, to find recurring activity sequences in the data (See STAR Methods). SeqNMF identified recurring neural activity patterns by solving the CNMF problem with an innovative regularization term that minimizes the redundancy of motifs. Specifically, SeqNMF tries to decompose a data matrix **X** ∈ ℝ ^*N* ×*T*^ into the sum of convolutions of *K* different motifs (factors) **W** ∈ ℝ^*N* ×*K*×*L*^ and their temporal occurrences **H** ∈ ℝ^*K*×*T*^, where *N* is the number of neurons and *T* is the total recording time bins (Fig 1D, see STAR Methods,^(37)^ for details). To apply SeqNMF to the trial-based imaging data^(29)^, we concatenated the Δ*F/F* calcium traces of all the trials into one large *N* × *T* matrix. Like other NMF methods, SeqNMF assumes the input data to be sparse and non-negative. These constraints are natural to calcium imaging data whose major components are sparsely distributed, non-negative calcium transients. We thus thresholded the data to keep the major calcium transients in the data to form a sparse non-negative input data matrix *X* (STAR Methods, Fig S2).

To ensure identification and validation of motifs potentially specific to certain trial types, we partitioned the data into training and test trials while balancing the number of different trial types within the training and test sets (Fig 1C, see STAR Methods for details). Though SeqNMF was able to identify some significant motifs **W** and their temporal occurrences **H** (Fig 1D), we observed two major deficiencies: First, the slow kinetics of calcium indicators resulted in ambiguous activation times of motifs, i.e., **H** is not sparse in time (Fig 1D), while the precise timings of motifs are crucial for the neural encoding of information (especially temporal coding). Second, the significant motifs found by SeqNMF algorithm are inconsistent between different runs of the algorithm with the same parameters, indicating that matrix decomposition by SeqNMF is sensitive to initialization when run on calcium traces (Fig 1E).

### SeqNMF optimization using Split Bregman Iterations

We speculated that the aforementioned problems with SeqNMF are, in fact, not caused by the model itself, i.e., the cost function being optimized, but rather by the model fitting procedure in SeqNMF which uses a multiplicative update rule^(38;39)^. To validate our belief, we revisit the SeqNMF cost function:

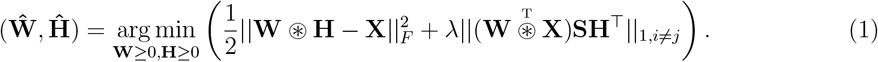

Here the first term is a reconstruction error term and the second term is a regularization term that reduces the redundancy of the motifs^(37)^. The reconstruction error term simply measures the difference between the reconstructed data 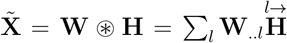 and **X**, where 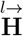 shifts a matrix **H** in the arrow direction by *l* time bins. The regularization term evaluates the overall cross-correlation (excluding self-correlation) between 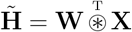 and **H**, with a smoothing matrix **S** ∈ ℝ ^*T* ×*T*^ (**S**_*ij*_ = 1 when |*i* − *j*| *< L* and otherwise **S**_*ij*_ = 0). 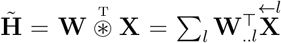 measure the overlap between motifs **W** and data **X**. Since the cost function is only convex in **W** and **H** separately but not jointly, SeqNMF alternate solves over **W** and **H** to find a local minimum:

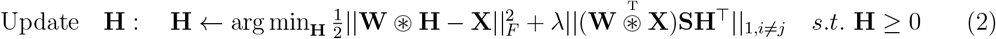

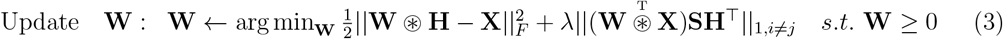

To solve Equations (2) and (3), SeqNMF adopts a multiplicative updating rule (MUR) inspired by the original NMF algorithm^(46;47;32)^, which ensures that **W** and **H** remain non-negative (STAR Methods). Prior works have shown that the MUR was susceptible to slow convergence, poor local minimum, and non-sparse solutions^(38;39)^. To alleviate these issues, we replace the MUR with a full solve for the optimization problems of Equations (2) and (3) using Split Bregman Iterations (SBI)^(45)^ (see STAR Methods).

The central idea of our SBI approach is to introduce an auxiliary set of variables **D** and **B** that represent the significant correlations between motif activations and the tolerance of solving the true optimization program, respectively. These variables are updated together with the actual variables of interest (e.g., the motifs or the activations), resulting in convergence to the solution of the original problem, but with greater robustness. For example, the SBI algorithm for solving Equation (2) iterates over the following three updates:

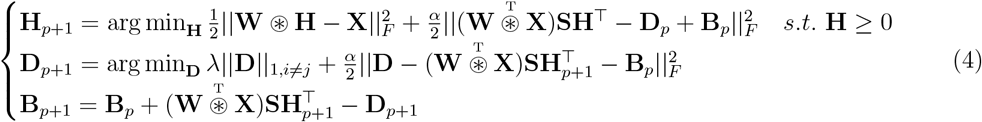

where **H** ∈ ℝ ^*K*×*T*^, **D** ∈ ℝ ^*K*×*K*^, **B** ∈ ℝ^*K*×*K*^, *p* is the iteration number. Note that as **B** → 0, **D** becomes a sparse approximation of the correlation term 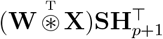 (i.e., the middle equation), allowing the LASSO-style regularization in Equation (2) to be replaced with a least-squares loss. This replacement greatly increases the algorithmic robustness. Equation (3) can be similarly solved by replacing **H** with **W** in the first line of (4) (See STAR Methods for details).

#### Hyperparameter selection

In SeqNMF, the hyperparameter *λ* balances the weights of reconstruction error and regularization error. A large *λ* will penalize the redundancy of motifs and produce a sparse factorization solution. Our new SBI-based updating rule introduces an additional hyperparameter *α*, which can be different when solving Equations (2) and (3). Thus there are two extra hyperparameters *α*_**H**_ and *α*_**W**_, corresponding to updates over **H** step and **W**, respectively. *α*, as seen in the first row of Equation (4), plays a similar role in balancing the reconstruction and regularization cost, indicating that *α*_**H**_ and *α*_**W**_ control the sparsity of **H** and **W**. To highlight the impact of *α*, we show how the same simulated data matrix factored with SBI can have non-sparse motifs **W** convolved with sparse temporal loadings **H**, and sparse motifs **W** convolved with non-sparse temporal loadings **H** (Fig. 2A). Notably, as we tuned *α*_**H**_ and *α*_**W**_, the original optimization cost in Equation (1) did not change, indicated that SBI was not indirectly changing the model with implicit regularization terms but simply enabling the specification of local minima of the global cost. In fact, adding regularization to the SeqNMF problem solved with MUR (e.g., ||**H**|| _1_ or ||**W**|| _1_) failed to produce similar sparse solutions as SBI (Fig. S4), suggesting that SBI is more powerful than MUR in solving the SeqNMF problem.

**Figure 2.**
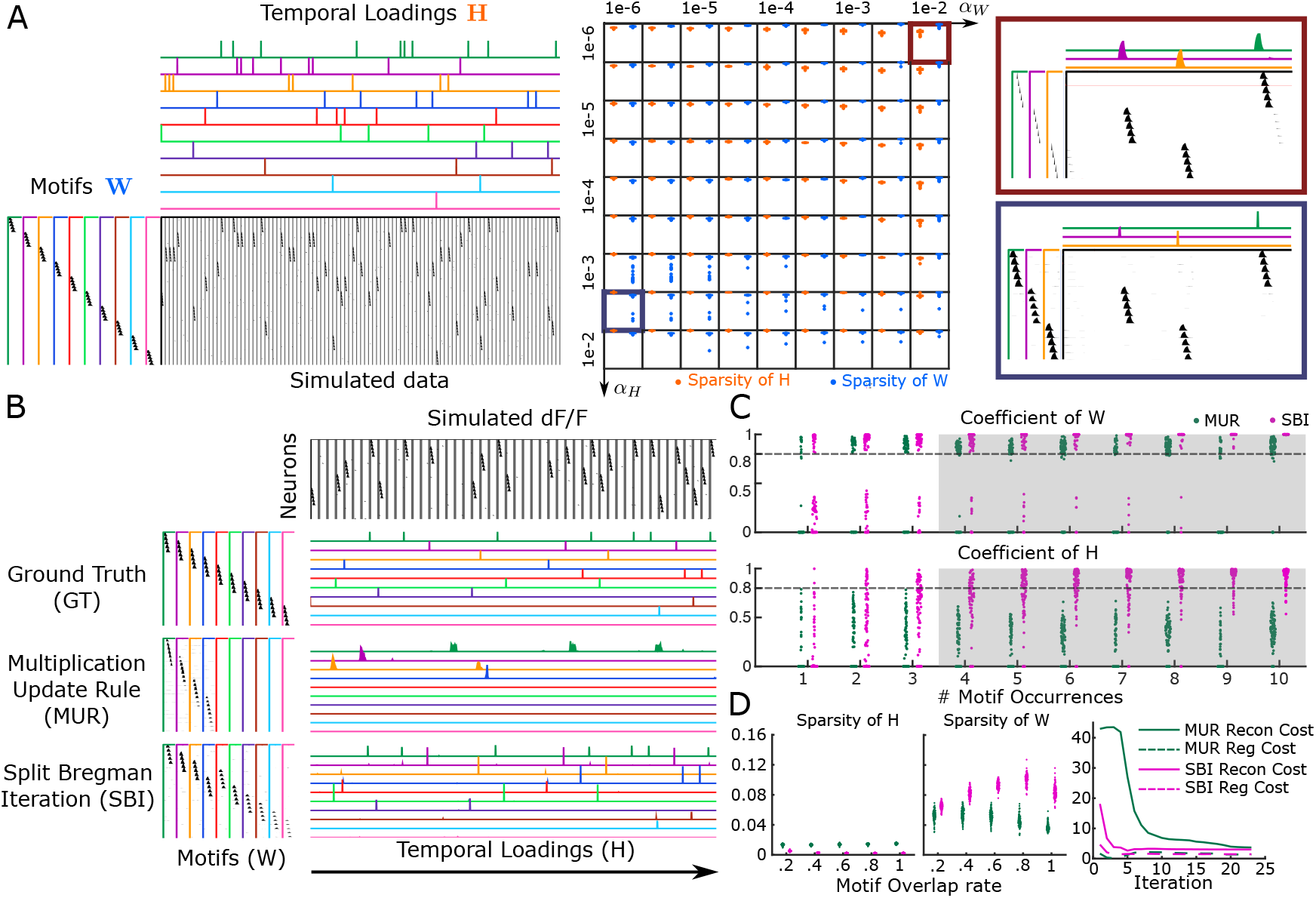
Split Bregman Iteration outperforms Multiplication Updating Rules on simulated data (A) Left: Example of simulated calcium imaging data. 10 different motifs occur different times (1 to 10) in the data. Right: Sparsity of solutions **H** and **W** as a function of hyperparameters *α*_**H**_ and *α*_**W**_ in Split Bregman Iteration (SBI). Tuning the hyperparameters *α*_**H**_ and *α*_**W**_ can drive SBI to produce solutions with non-sparse **H** and sparse **W** (top right panel) or sparse **H** and non-sparse **W** (top left panel) (B) Comparison of factorization results of MUR and SBI against ground truth when motifs have shared neurons. From top to bottom rows: simulated dF/F traces, ground truth motifs **W** and their temporal loadings **H, W** and **H** found by MUR and SBI. (C) Coefficients of **W** and **H** to the ground truth over 100 runs of MUR and SBI on simulated data. Coefficients increased as motif occurred more times in the data. SBI can identify motifs with high fidelity when the motif occurres more than 4 times (shaded grey region). (D) Left: Sparsity of **W** and **H** as a function of motif overlap ratio. SBI produced motifs with greater sparsity in their temporal loadings **H** while preserving more activities in motifs themselves **W** than MUR. Right: SBI converges faster than MUR.

### SBI motif detection approach robustly finds temporally precise motifs

The improved robustness and control of sparsity enabled by SBI allowed us to further study motifs in auditory cortex. Specifically, in the auditory cortex, the precise timing of neural activity is crucial for understanding the function of neural circuits since sounds are rapidly time varying stimuli. Thus we employed SBI to steer SeqNMF to produce sparse temporal loadings **H** and non-sparse motifs **W** by increasing *α*_**H**_. Simulation-based validation in synthetic data using the optimal choice of *λ* in SeqNMF (STAR Methods, Fig. S6) verified that SBI could identify motifs **H** with more precise timings and greater similarity to the ground truth than MUR (Fig. 2B,C). The SBI solutions bear additional advantages over MUR solutions, including greater convergence speed (fewer iterations to converge) and robustness to initialization and noise (Fig. 2D). These characteristics hold regardless of the noise type and noise level being added to the data (Fig. S3A, S3B, S3C, S3D).

#### Assessing significant motifs

One final challenge in motif detection is in determining when identified motifs accurately reflect data statistics, e.g., because motifs may present a limited number of times, posing uncertainty to successful identification. We thus ran additional benchmarks to identify the minimum times the motif should present in the data to be reliably detected (see STAR Methods). We found that, under reasonable noise levels, SBI-based seqNMF was able to robustly identify motifs as significant when they appeared in the data as little as four times, while the MUR-based algorithm required many more repetitions to faithfully find them (Fig. 2C). These results were consistent across noise levels and noise types (Fig. S3A, S3B, S3C, S3D), validating SBI as an improved algorithm for motif discovery.

### SBI precisely identified motifs on 2-photon imaging data on A1

Following our synthetic-data validation, we applied our SBI to the published two-photon imaging dataset to test our initial hypothesis that motifs in A1 encode both stimulus and choice. To ensure that each analyzed session included enough active neurons (*N >* 10) and trials (*Tr >* 10 for each frequency), we restricted our analysis to 24 sessions fulfilling these criteria (“good sessions”)(STAR Methods, Table S1). To run SBI we fit the three hyperparameters (*λ, α*_**H**_ and *α*_**W**_), via a semi-automatic procedure (see STAR Methods). At a high level, the procedure aimed to find the hyperparameters under which most ‘motif’ structures in the data are captured while the temporal precision (sparseness) of the motifs are preserved. We validated our procedure by plotting the temporal sparseness and the proportion of captured motifs across different *λ* and *α*_**H**_ (STAR Methods). The optimal hyperparameter settings, despite including additional parameters, coincided with the optimal point in the SeqNMF paper^(37)^ where changes in reconstruction cost and changes in regularization cost are balanced (Fig. S7).

We applied the hyperparameter selection procedure to the training data of all the 24 good sessions (Table S3). Once the motifs were extracted on training data, we used them to fit the remaining test data (Fig. 3A, STAR Methods). We thus acquired the accurate occurrence times of all the motifs in each session. Based on our simulation-based analysis of motif significance in SBI we dropped any motif that appeared *<*4 times. To further interpret the motifs, we developed statistical tests to identify neurons that are significantly activated within each motif (STAR Methods). Beyond the statistical tests we also validated each significant motif and their active neurons by visualizing the neural activity relative to each occurrence of the motif (Fig 3B).

**Figure 3.**
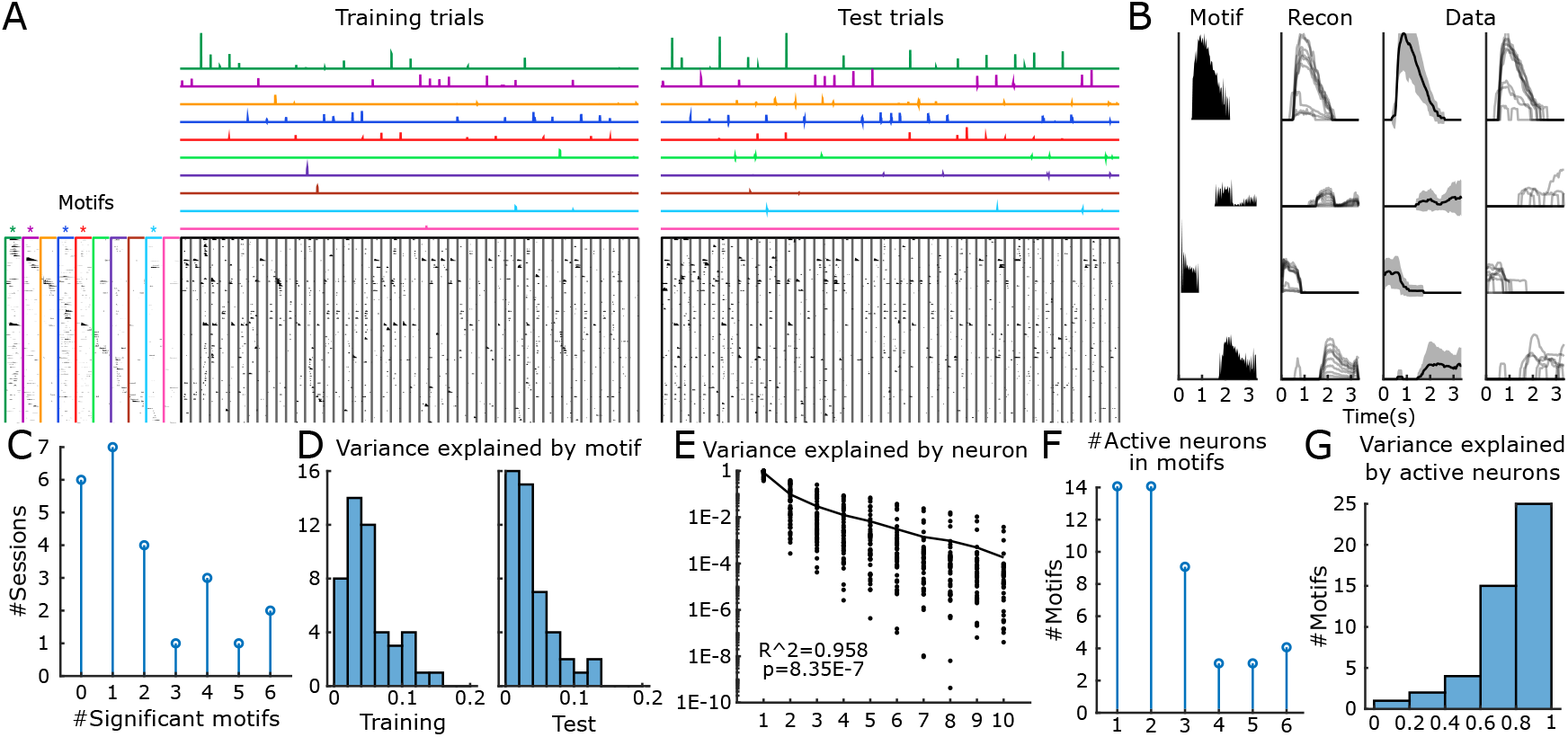
General statistics of motifs captured by new algorithm (A) Motifs (left) and their temporal occurrences (top rows) in the training and test trials detected by SBI approach in one example session. Significant motifs were labeled with an asterisk on their top. (B) Validation of the significant motifs and the active neurons. Each row represent one neuron. Left panel: the motif itself. Second panel: Reconstruction of neuralk activities from motifs and their temporal loadings. Third panel: mean (black lines) and standard deviation (shaded regions) of dF/F traces over the time windows of all the occurrences of the motif. Right panel: dF/F traces over the time window of each motif occurrence. (C) Histogram of the number of significant motifs in each session. (D) Histogram of the variance explained by the significant motifs in training data (left) and test data (right). (E) The proportion of variance explained by the active neurons within the motifs, sorted in descending order. The variance decay quickly with an exponential rate, indicating that only a small group of neurons captures most activities of the motif. A linear model was fit to the logarithm of the neuron’s explained variance and its index (*R*^2^ = 0.958, *p* = 8.35 ∗ 10^*−*7^). (F) Number of active neurons in significant motifs. (G) Variance explained by all the active neurons within motifs.

Among the 24 good sessions, the algorithm captured an average of 1.96 significant motifs per session (95% confidence interval: 1.15-2.77, Fig. 3C). Each significant motif captured 5.03% of the total neural activity variance in the training data (95% confidence interval: 3.63%-6.43%) and 3.64% in the test data (95% confidence interval: 2.26%-5.02%, Fig. 3D, STAR Methods), suggesting that a small but significant fraction of neural activities are structured as neural motifs.

Studying the neural motifs (each a neuron× -motif duration matrix) configurations revealed that a small group of neurons capture the major proportion of motif variance, and the numbers of significantly active neurons whin motifs are small(Fig 3E,F,G *p* = 8.35 ∗ 10^−7^, STAR Methods).

We next investigated if the active neurons within motifs exhibit any specific spatial distribution pattern (e.g., if they are clustered or regularly ordered in space) in the imaging field-of-view (FOV, 370 × 370 *µm*). We used Clark-Evans R index to quantify the clustering of motif neurons^(48)^ and tested against the null hypothesis that the neurons are completely randomly distributed using bootstrap (STAR Methods). *R <* 1 indicates spacial clustering, while *R >* 1 indicates regularly spacing. To ensure statistical power, we only tested on motifs with at least 5 active neurons. Among all these motifs, the spatial pattern of active neurons are indistinguishable from complete spatial randomness (CSR). However, given the small number of active neurons, it is still possible that our tests failed to capture their special spatial patterns (clustering or regular spacing). Together, the above statistics suggested that motifs involve a small group of active neurons that are likely to be randomly distributed in space.

### A1 motifs encode stimulus or stimulus and choice jointly

The precise motif occurrence times provided by SBI (STAR Methods, Fig S8A) revealed that many motifs exhibited selectivity to certain trial types (Fig. 4A). To quantify this occurrence selectivity, we applied an encoding index ranging from 0 to 1 for each task variable or combination of variables (STAR Methods). Given that we were analyzing motifs in A1, we expected to observe motifs with greater selectivity to auditory information than non-auditory information. Unsurprisingly, the significant motifs in A1 shows greater stimulus and joint encoding indices than choice and lick encoding indices (Fig. 4B). Note that stimulus and choice are highly correlated in each session and the number of trials of certain type is at times limited (Table S1), posing challenges in disambiguating between the motifs’ encoding of different variables. We thus developed a procedure based on multiple Fisher’s exact tests conditioned on certain trial types to disentangle stimulus-encoding and choice-encoding (STAR Methods). Following our tests, we found a large number of motifs encoding stimulus, and a small amount of motifs encoding stimulus and choice jointly (Fig. 4C, top). Interestingly, we did not find motifs that are strictly choice-encoding or lick-encoding (see Table S4 for complete encoding statistics).

**Figure 4.**
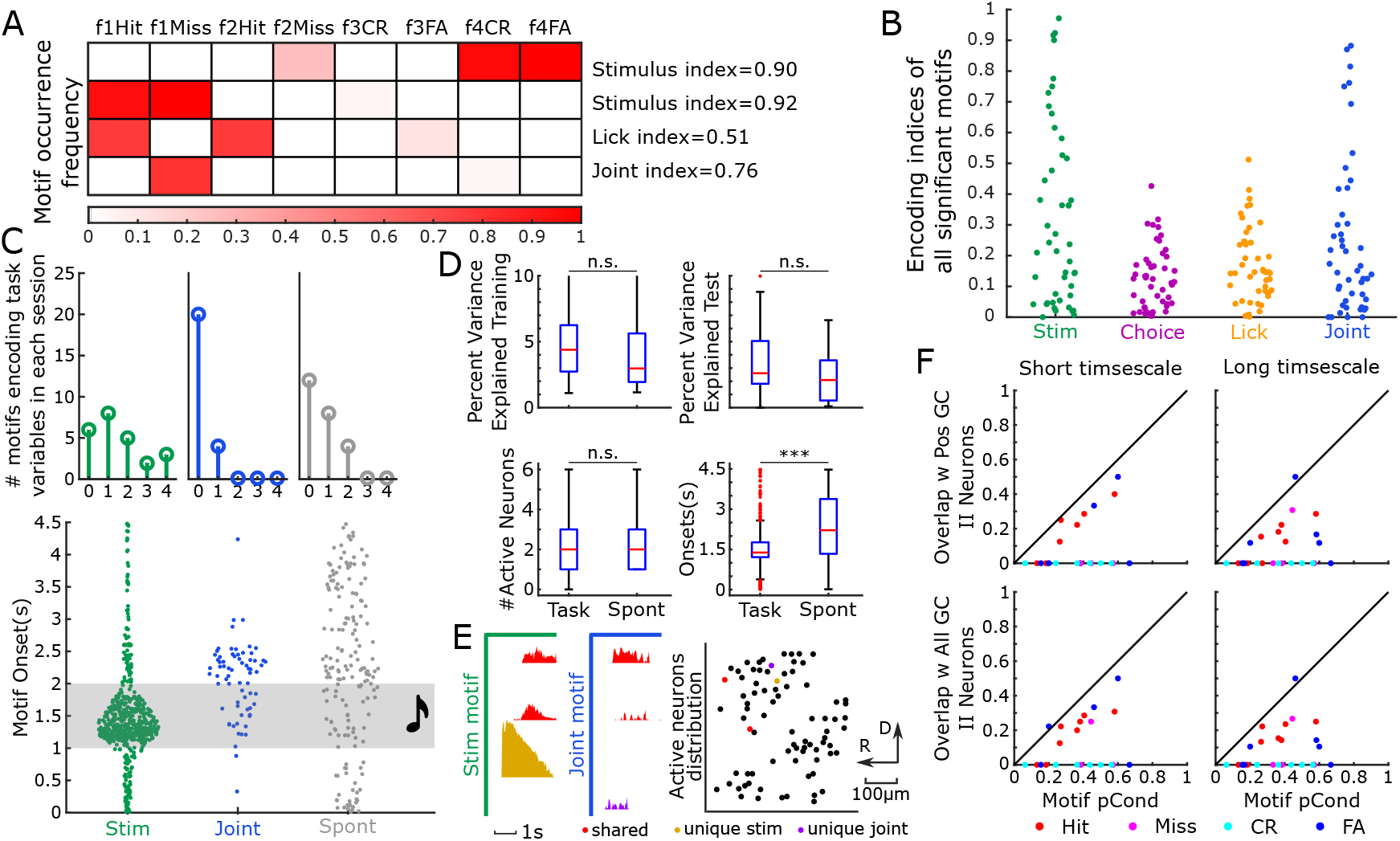
Motifs encode various task variables in a sound discrimination task (A) Examples of motifs preferring to occur in certain trials. Each row is one motif, the heatmap indicates the conditional probability of the motif occurrence in each type of trials. Each motif’s encoding index for specific task variables is labeled on the right. (B) Distribution of the encoding indices of different task variables in all the significant motifs. (C) Top: Histogram of the number of task-encoding motifs and spontaneous motifs in all the sessions. Bottom: Histogram of the onset time of the task-encoding motifs and spontaneous motifs in all the sessions. Trials started at 0s, the tones were played during 1-2s (shaded region). (D) Top left: the variance explained by the task-encoding motifs in the training data is not significantly different from the spontaneous motifs(two-sampled t-test, p=0.1839). Top right: same test on the test data(two-sampled t-test, p=0.0825). Bottom left: the number of active neurons is not significantly different between task-encoding motifs and spontaneous motifs(two-sampled t-test, p=0.6154). Bottom right: the variance of onsets of task-encoding motifs is significantly smaller than spontaneous motifs(*p* = 1.5048 ∗ 10^*−*22^, two-sample one-sided F-test). (E) An example of two different task-encoding motifs with shared neurons. The stimulus-encoding motif (left, green) encodes f1 while the joint encoding motif (middle, blue) encodes f1 and miss jointly. Two neurons (red) are active in both motifs, while each motif contains an unique active neuron. Right: spatial distribution of these active neurons in the field of view. Black dots are all the other neurons. (F) The overlap of the active neurons in the task-encoding motifs and the GC-linked intersection information(II) neurons in Francis et al. GC links were estimated at short(233ms, left) and long(1033ms, right) timescales. Positive GC links only (top) and all GC links (bottom) were selected to compute the overlap, measured as the dice coefficient.

We next investigated the motif onset times within trials. Most stimulus-encoding motifs occur during sounds, while the joint-encoding motifs tend to start later than stimulus-encoding motifs (Fig. 4C bottom), indicating the cortical processing of sensory information to drive behavior. We were surprised to find that some sound-encoding or joint-encoding motifs start earlier than sound onsets, which should not be possible since both the next trial’s sound category and the intertrial interval are random and as such the prediction of the next trial is impossible. By inspecting the original fluorescence(dF/F) traces of motif neurons, we found spontaneously active neurons in the motifs. Specifically, these neurons were often suppressed at the tone onset, however certain spontaneously active neurons tended to precede specific motifs. These regularities were captured by the motif identification, indicating that while there is no information of the upcoming tone, the way the neural population responds to a stimulus can be dependent on the spontaneous activity right before the stimulus (Fig S9).

In addition to task-encoding motifs, we also found spontaneous motifs that do not encode any captured task variable (Fig. 4C top). Contrary to the task-encoding motifs whose onsets are narrowly distributed within trial, the onsets of spontaneous motifs exhibit a significantly wider distribution (*p* = 1.5048 ∗ 10^−22^, two-sample one-sided F-test), while other statistics between task-encoding motifs and spontaneous motifs are similar (Fig. 4D top). Further studies could explore the potential roles of these spontaneous activities during and independent of a task, e.g., representing body movements or other internal signals.

Although randomly distributed in space, active neurons can be shared between different task encoding motifs. For instance, some active neurons are shared between a stimulus-encoding motif and another joint-encoding motif that encodes the same stimulus and a specific choice (Fig. 4E), while each encoding motif has unique active neurons. Such shared representations align with concurrent population coding theories^(49)^. Taken together, our motif characterization verifies the existence of neural motifs encoding sensory and behavioral information in A1.

### Motif neurons partially overlap with GC-linked neurons

Prior work demonstrated that A1 contains a subset of neurons that transiently carry intersection information(II), i.e., the sensory information used by the neurons to inform behavior choices^(29;50;51)^. Specifically, a small group of II neurons formed functional subnetworks through sparse Granger causality(GC) links. As a strong GC link indicates that the recent history of a neuron can improve the prediction of another neuron’s activity^(52;30;29)^, we speculated that the GC-linked neurons in A1 should overlap with the active neurons in our task-encoding motifs. Since the GC analysis were performed separately on the hit, miss, false alarm (FA) and correct rejection (CR) trials, we grouped the motifs by their highest conditional probability of occurrence in these four trial types (STAR Methods). The overlap between neurons active in motifs and GC-linked neurons (quantified by the dice coefficient) increased with the conditional probability of occurrences (Fig. 4E). The fact that the points in Figure 4E fall below the *y* = *x* line indicates that some motifs can have high conditional probabilities in certain trial types but low overlap with II neurons, while motifs never have a high overlap with II neurons without also having high conditional probabilities. This divergence could be explained by two factors: First, we thresholded the fluorescence (dF/F) traces and only kept super-threshold transients before finding motifs, while GC analysis used the entire fluorescence traces; Second, sequential activation of neurons (motifs) does not necessarily prove causality, while causality is sufficient to guarantee sequential activation of neurons.

### Additional 2AFC sound discrimination task verifies the prevalence of stimulus and joint-encoding motifs

Given that the animal was performing a go/no-go task, the joint-encoding motifs we identified so far may also be interpreted as motifs representing the stimulus and the animal’s engagement states rather than decisions. Thus, to test the generalizability of our observations of both stimulus and choice encoding in A1, we trained additional mice (8 mice, 40 sessions) to discriminate different sounds in a 2AFC task, by turning a wheel (Fig. 5A). Specifically, the animals learned to turn a wheel in one direction in response to low-frequency sounds and in the opposite direction in response to high-frequency sounds^(53)^ (Fig. 5A). Water reward was delivered immediately after the animal made the correct wheel turn. This new behavior paradigm thus distinguishes decision, engagement, and reward in each trial and allows us to explore the relationship between motifs with other task variables such as movement and reward. In these mice we imaged areas of auditory cortex (A1 and the dorsomedial field, DM) and then performed the same motif analysis on the two-photon imaging data as before. We included both A1 and dorsomedial field (DM) in our analysis because DM was previously thought to be part of A1 but might also be considered an independent region^(54)^. By imaging a larger FOV (500*µm* × 500*µm*) and using Suite2P to identify cells automatically (STAR Methods), we obtained 310 neurons on average (compared to 41 neurons in the published dataset with a 370*µm* × 370*µm* FOV), thus enabling motif identification within a larger neuron population.

**Figure 5.**
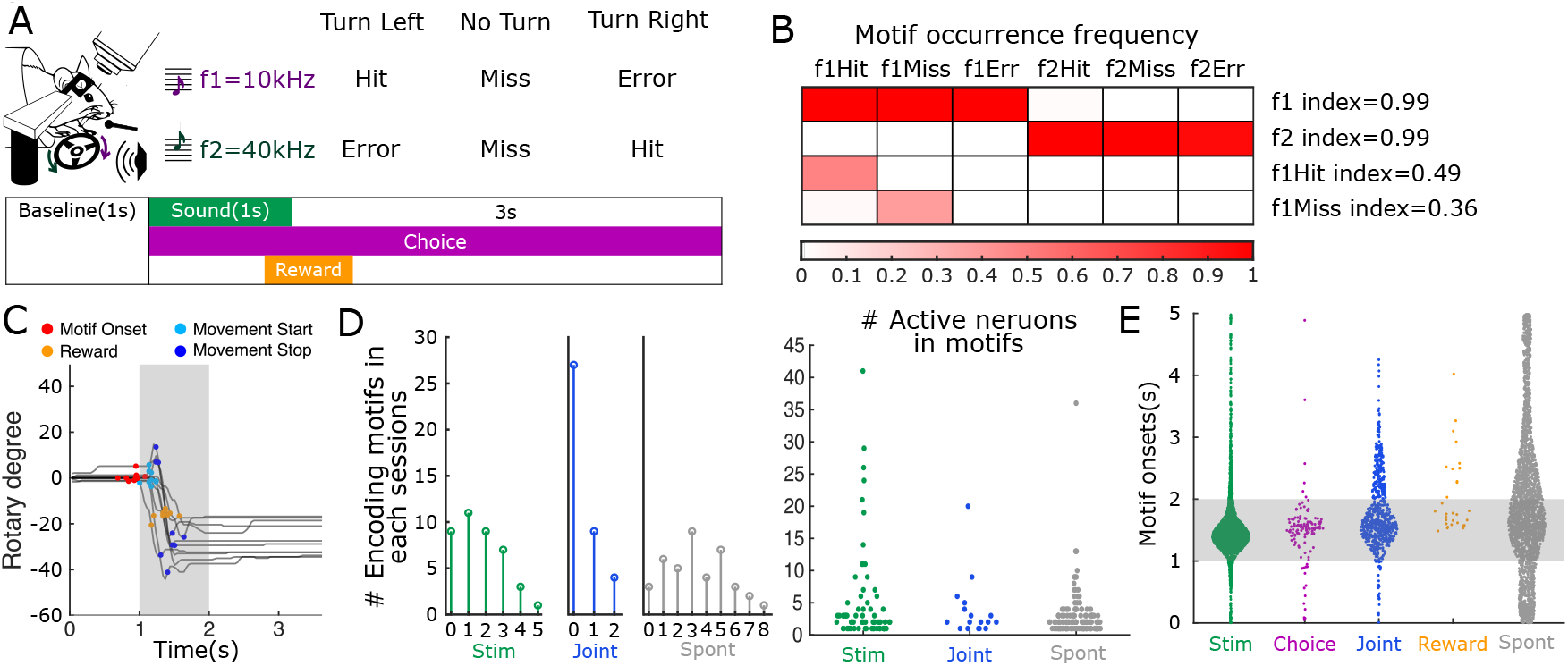
Motifs encode more variables in an additional 2AFC sound discrimination task (A) Task diagram: mice were trained to turn a wheel to the direction associated with sound frequency during the choice window for a water reward. (B) A few example of task-encoding motifs, similar to Fig 4A. (C) Wheel rotary trajectories of all the trials in which one motif occur. The timings of movement start, movement stop, reward, and motif onsets are labeled in each trial with different colors (See STAR Methods). (D) Left: Similar to Fig 4C, histogram of number of different motifs in all the sessions; right: histogram of the number of active neurons in different motifs in all the sessions. (E) Similar to Fig 4C, histogram of the onset time of the task-encoding motifs and spontaneous motifs in all the sessions.

Consistent with our findings above, we discovered widespread stimulus-encoding motifs and a small number of joint-encoding motifs (Fig. 5B,D, Table S5). The joint-encoding motifs we identified are mostly active in correct and miss trials, with only one motif encoding high frequency and error trial (Fig. S11A), suggesting that both choice and disengagement can be jointly represented with stimulus by neural motifs. We also identified very few purely choice-encoding motifs (n=2), which were not observed in the published dataset, probably due to the smaller neuron population being analyzed.

Given that we imaged A1, it was expected that we would encounter stimulus-encoding motifs in almost all the sessions. However, joint-encoding motifs were only observed in about 1/3 of the sessions. To explain this inconsistency among sessions, we tested several hypotheses, including the effect of imaging FOV, imaging depth, individual variability, the number of active neurons and the degree of balance between different trial types. The number of joint-encoding motifs in different sessions can not be explained by the difference in imaging FOV (two-sample t-test between DM and A1, p=0.87), imaging depth (linear regression model, F-stats=0.0177, p=0.895), and the number of active neurons (linear regression model, F-stats=3.55, p=0.067). However, joint-encoding motif count varies by individual (chi-square test, *χ*^2^ = 17, p=0.0175), and is significantly affected by the balancing index between different trial types (i.e., more motifs being found when there are more miss or error trials, see STAR Methods). These results indicate that joint-encoding motifs exhibit individual variability and are more likely to be found when there are more balanced trial types.

To further explore motifs potentially associated with other task variables, we tested if the motif onsets were time-aligned to a number of task-related events, including wheel movement onsets and offsets as well as consumption of water rewards (Fig. 5C, STAR Methods). This analysis identified reward encoding motifs (n=2 in one session), but did not identify any motif whose onsets are aligned with wheel movement onsets or offsets. In comparison, we did not discover reward-aligned motifs in the published data, where licking was used for both decision and reward (Table S4), potentially due to the inability to disambiguate between the co-occurrence of joint-encoding motifs and reward-encoding motifs.

Similar to the published dataset, the motifs we found during our 2AFC task take up a small proportion of total neural activity variance and consist of small number of randomly distributed active neurons (Fig. 5D, Fig. S11B), suggesting that task structure has little influence on motif neuron statistics. Their onsets also show similar distributions as the published dataset (Fig. 5E), suggesting that the task structure has little influence on the motif onsets. Together, the similarity of the results from the different task designs suggests that the occurrence of stimulus and joint motifs are independent of task structure but that the occurrence of reward motifs might depend on the separability of the decision and reward phase.

### Mixed and multiplexed encoding of task variables by single neurons within motifs

So far, we identified neuronal motifs, i.e., groups of cells activating together in a particular pattern. We next wanted to investigate the composition of the motifs. Do single neurons within motifs show similar or heterogeneous encoding properties? Single neuron in A1 can exhibit diverse auditory or non-auditory encoding properties^(21;25;30)^. We thus investigated the encoding properties of single neurons, including the encoding of stimulus, choice, stimulus & choice, reward, wheel movement, and compared them against the encoding properties of motifs. Overall we want to test between two hypotheses: the encoding properties of motifs might be simply the aggregation of neurons with the same encoding properties (Hypothesis I, Fig. 6A left), or the motifs consist of single neurons with misaligned encoding properties, i.e. population codes^(55;49)^ (Hypothesis II, Fig. 6A right). To disentangle stimulus, choice and reward information, we applied a testing procedure similar to testing motif encoding. Since neurons in the auditory cortex can be responsive to sound onsets or offsets^(56)^, we defined stimulus-encoding neurons by comparing the neuron’s activity level against baseline in three different 0.5s time windows during sound or after sound. The neuron was defined as encoding the sound frequency if its activity was greater than the baseline in any of the windows (STAR Methods). For choice encoding, we tested if the neuron’s post-sound fluorescence responses significantly differ between trials of different choices conditioned on a specific stimulus (STAR Methods, Fig. S12C). For reward encoding, we tested if the calcium transient times of each neuron are time-aligned to reward events, conditioned on trials with a certain stimulus. Additionally, given that there can be spontaneous movements before trail onsets, we also tested the encoding of wheel movement for each neuron by evaluating the cross-correlation between wheel turning speed and neural activity during the inter-trial intervals (STAR Methods, Fig. S12D).

**Figure 6.**
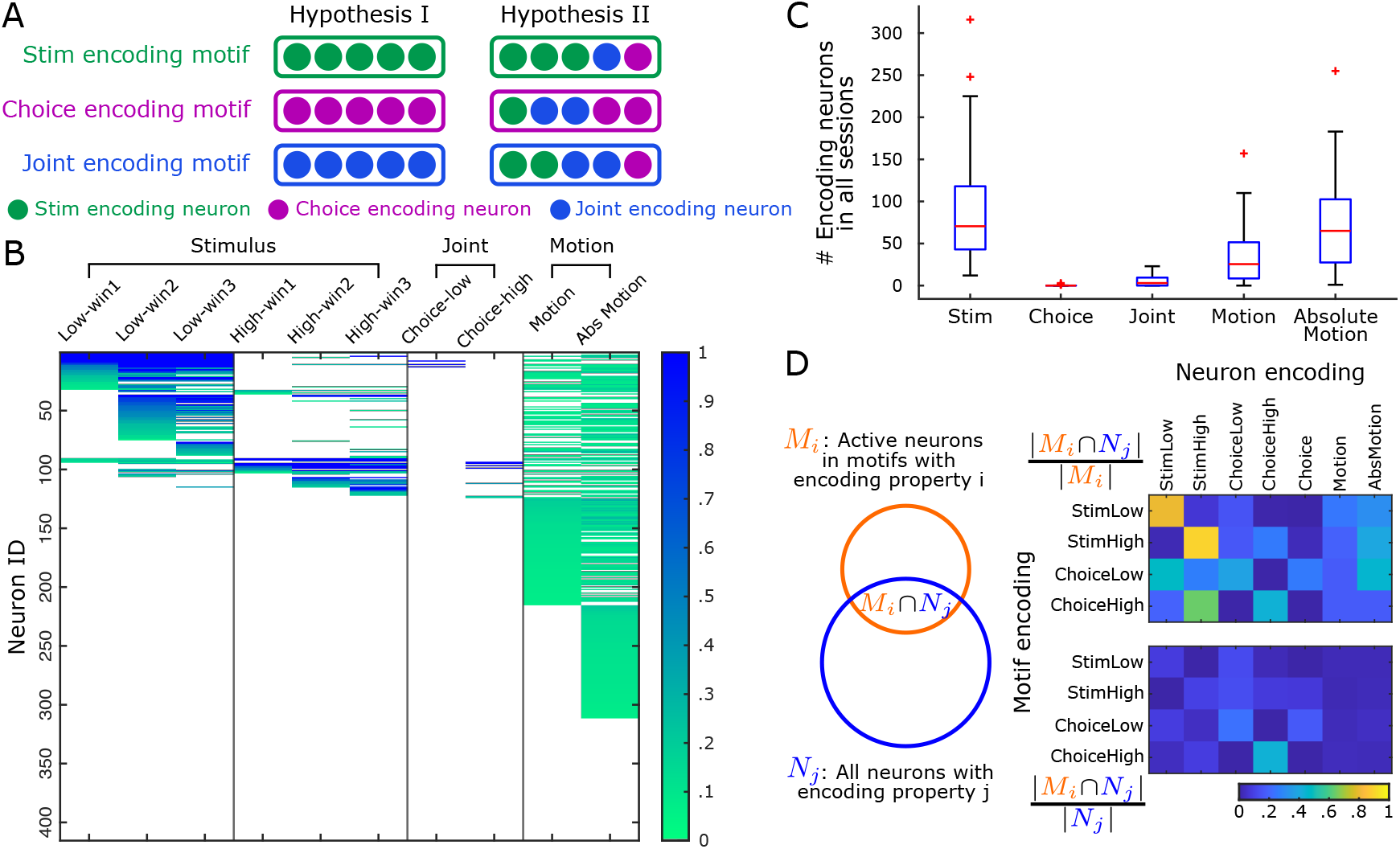
Mixed and multiplexed encoding properties of single neurons within motifs (A) Diagram of two hypothesis we want to test. Left: motifs are consist of single neurons with the same encoding properties. Right: motifs are consist of neurons with mixed encoding properties (population code). (B) Encoding of different task variables by single neurons in one session. Each column is all neurons’ encoding for one variable (within a certain time window). All colored grids indicate significant encoding. In the Stimulus and Joint columns, grid color represents the difference in the mean dF/F activity between a certain time window against baseline, indicating encoding strength. The neurons that are significantly activated in any of the three 0.5s time windows related to a certain sound (i.e., colored in any of the first three columns or any of the 4th to 6th columns) are defined as neurons encoding the corresponding sound frequency. In the last two columns, grid color represents the maximum cross correlation between neural dF/F and wheel movement speed or absolute wheel movement speed, indicating the strength of movement encoding (STAR Methods). (C) Number of different encoding neurons across all sessions. (D) Left: schematic diagram of how we compared the motif encoding properties and single neuron encoding properties. Top: the proportion of the overlap between the active neurons of motif with encoding property i (indicated by row name) and all neurons with encoding property j (indicated by column) over all active neurons within motif. Bottom: the proportion of the overlap between active neurons with encoding property i (indicated by row name) and all neurons with encoding property j (indicated by column) over all neurons with encoding property j (See STAR Methods).

Consistent with concurrent studies based on large-scale neural recordings^(26;21;57)^, we observed diverse and heterogeneous representations of task variables by single neurons. We discovered substantial stimulus and wheel movement encoding neurons in each session (Fig. 6B). The movement-encoding neurons show weak but significant cross-correlations to wheel movement. Additionally, there are a small number of joint-encoding neurons, very few purely choice-encoding neurons, and no reward-encoding neurons(Fig. 6C).

To test between our two hypothesis, we compared encoding properties of motifs and their active neurons. To do this, denote the set of active neurons of a motif with encoding property i as *M*_*i*_, denote the set of all the neurons with encoding property j as *N*_*j*_, we computed two ratios: the overlap over all motif neurons 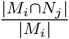 and the overlap over all encoding neurons 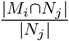. When I and j are the same, the first ratio measures the proportion of active neurons that share the same encoding property as the motif they belong to, the second ratio measures the proportion of neurons that are active within a motif over the entire neuron population with the same encoding property. We plotted the two ratios over all the motifs (take the mean over all motifs) for all the combinations of motif encoding property i and neural encoding property j in Fig. 6D top and bottom. If the first hypotheses holds, we would see diagonal structure in both plots, and the diagonal part should be close to 1 in the first plot. We found that motifs actually consist of neurons with mixed encoding properties, and these properties do not always align with the motifs’ encoding properties (Fig. 6D top). Specifically, stimulus-encoding motifs (top two rows) are mostly consist of neurons encoding the same frequency, while joint-encoding motifs (bottom two rows) often include neurons with more heterogeneous encoding properties. The bottom plot illustrates over the entire neuron population with a certain encoding property, what proportion of neurons belong to a motif with a certain encoding property. We found that all the elements were small (*<* 10%), except that a slightly higher proportion of joint-encoding neurons are active in joint-encoding motifs (*>* 20%), suggesting that joint-encoding motifs are more likely to recruit joint-encoding neurons. Moreover, neurons can be shared between different motifs, with the highest proportion being shared between motifs with similar encoding properties (e.g. high-frequency vs high-frequency, or high-frequency vs highfrequency and correct choice jointly) (Fig. S13). Despite very rare wheel movement encoding motifs and substantial movement-encoding neurons, there are always a small proportion of movement-encoding neurons participating in all types of motifs. Taken together, our study indicates the complex mapping between motifs and single neurons, in which neurons are reused by multiple different motifs (multiplexed encoding), while multiple neurons with diverse encoding properties constitute one single motif (mixed encoding). This mismatch between neuron and motif encoding properties reveals that, as a sparse population code^(55;49)^, motifs serve as a parallel encoding scheme in transforming sensory information into behaviors.

## 3 Discussion

To date, the language of cortical function has traditionally focused on the lens of characterizing single neuronal encoding. However, neural populations project downstream and across brain areas as both individuals and populations, begging the question of how different pieces of information can be read out from the same group of neurons. By exploring the tuning properties of both single neurons and spatio-temporal motifs, we provide evidence that different levels of neural population codes can contain distinct information, allowing other brain areas to multi-plex the information by being sensitive to either individual units, or the population in total.

Specifically, we study here such multi-scale encoding in the context of mouse Auditory Cortex during a decision making task. We find that at the neuron population level, neuronal motifs, which have been considered as a fundamental population code for cognition can encode a sound stimulus, jointly encode a stimulus and a choice, or occur spontaneously. The motifs comprise a small set of active neurons with mixed encoding properties inconsistent with the motif’s encoding properties, indicating that information is represented differently by single neurons and neuron population. Our results thus reveal two separate encoding schemes at different scales in the auditory cortex.

This study was made possible by the development of a new approach to identify neuronal motifs with high temporal precision in calcium imaging. We validated our new algorithm with synthetic data to show its robustness to different noise. Our method is not restricted to auditory cortex and can be applied to identify more motifs in other brain regions or across multiple brain regions, enabling us to investigate the fundamental role of motifs multi-resolution neural encoding and in cognition.

### Precise motif identification on calcium imaging data

Neural motifs play important roles in cortical information processing. To robustly identify neural motifs and to address current challenges that motifs lack temporal resolution due to the wide temporal response kernel in calcium imaging, we devised a new updating rule based on Split Bregman Iteration^(45)^ for the optimization program in SeqNMF^(37)^. Our new approach detects motifs with greater fidelity, requires fewer motif occurrences in the data, and converges faster than SeqNMF. We demonstrated the capability and robustness of our algorithm on simulated data under various noise conditions, and determined that a motif need to occur ≥ 4 times in the data to be faithfully reconstructed (Fig 2, S3A, S3B, S3C, S3D). Our algorithm managed to discover motifs and their precise activation times in two datasets, demonstrating the generalizability of our method in calcium imaging data.

### Sparse representation of sound and behavior information by neural motifs in A1

We observed that each significant motif only accounts for a small proportion of total neural activity (around 4%) and consists of a small group of spatially randomly distributed active neurons (Fig 3). The number of significant motifs within each session is also limited (≤ 5 in most sessions), indicating that most neural activity in A1 is not stereotypically structured. Nevertheless, many identified motifs were associated with a certain sound, a sound and a choice jointly, suggesting they may play important roles in sensory processing and cognition. Here we propose that most neurons are actually engaging in different motifs at various spatial scales, and we only observe a small proportion of them due to the following hypothesis. First, the active neurons within one motif can be located far away from each other, such as in different layers of A1, different subfield of auditory cortex, or even in different brain regions (such as in auditory cortex, parietal cortex and motor cortex), making them unlikely to be captured in our limited field of view (FOV) in A1 L2/3. The bottom-up projections from A1 L4 to L2/3 and rich modulation signals L2/3 receive from other brain regions have indirectly supported this hypothesis^(25)^. Second, given the large number of spontaneous motifs we observed, it is possible that many of the neurons are part of other motifs unrelated to our sound discrimination tasks. These motifs could be driven by unobserved behavioral and cognitive states ^(28;58)^. Future studies using larger scale methods for longer duration could explore these possibilities.

Additionally, we noticed heterogeneous occurrence frequencies between different types of neural motifs, with purely sound-encoding motifs being most frequent, joint-encoding motifs presenting in about 1/3 of the sessions, and other task-encoding motifs being very rare. This is consistent with our hypothesis that neurons across multiple brain regions constitute non-auditory encoding motifs, as these non-auditory motifs are less likely to be detected in a single FOV of auditory cortex.

### Motif analysis cannot be regarded as causality test between neurons

In A1, a small group of neurons transiently carry intersection information(II) and form sparse Granger causality(GC) networks^(30;29)^. As GC links indicate the existence of motifs, we compared the task-encoding motif neurons against the GC-linked II neurons. We observed that their overlap (measured as dice coefficient) is less than or equal to the conditional probability of the presence of the motif within a certain trial type. Theoretically, if a motif has a conditional probability of 1 in one trial type, the active neurons involved in that motif should completely overlap with the GC-linked neurons in that trial type. But since no such perfect motif exclusively occurred in one trial type, plus our pre-processing procedure dropped off less responsive neurons and under-threshold calcium activities, there is great heterogeneity in the overlap between the two neuron populations. Another information gap could arise from the lack of a causality test when detecting motifs, resulting in neurons that are sequentially activated, but are not causally linked. This comparison indicates that motif analysis cannot be regarded as a causality test between neurons.

### Potential roles and origins of task-encoding motifs

Neural motifs have been considered as fundamental building blocks of sensory processing and cognition ^(8)^. When mice were engaged in a sound discrimination task, stimulus-encoding motifs are mostly time-aligned with sound onsets. In contrast, behavior-related motifs begin in the later phase of the trials, indicating that purely sensory-encoding motifs may reflect local computations for sensory processing, while behavior-encoding motifs reflect global computations involving other brain regions. Since L2/3 of the auditory cortex receives rich top-down inputs or inter-cortical inputs, we speculate that the non-auditory encoding motifs we found could reflect the integration of local representation of sensory information in A1 and modulation signals elsewhere^(25)^. The modulation signal can arise from orbitofrontal cortex (OFC)^(59;60)^, posterior parietal cortex (PPC)^(61)^, second motor cortex (M2)^(62)^ or anterior cingulate cortex (ACC)^(63)^. The top-down projections from PPC or OFC can carry important information about current brain states and can reshape sensory cortical representations of external stimuli^(64)^, while movement modulation signals from M2 or ACC may carry important expectation signals, making the animal adaptive to the motor behavior in the task^(25;21;65)^.

### Motifs and single neurons are parallel encoding schemes on different scales

We observed substantial stimulus-encoding and movement-encoding neurons, a small amount of joint-encoding neurons, and very occasional choice-encoding neurons (Fig. 6C), which is consistent with the current idea of ‘Everything Everywhere’^(26;57)^. Nevertheless, individual neurons exhibit mixed encoding properties that are inconsistent with the motif encoding properties. Population encoding theory proposes that neurons with mixed encoding properties together can represent different information^(55;49)^, and these mixed encoding properties of motif neurons can contribute to an enhanced information encoding capacity of neuron ensembles^(66;67;68)^. The same neurons can be reused by different motifs, with most neurons being shared between motifs encoding the same stimulus. Such a multiplexed encoding property suggests that motifs serve as flexible building blocks for cognition and can dynamically change their compositions under different contexts^(69)^. Additionally, despite substantial wheel movement-encoding neurons, very few of these neurons form motifs time-aligned to movement onsets or offsets, indicating that most motion signals in the auditory cortex belong to modulation effects at the single neuron level instead of neuron population level. Among the entire neuron population with a certain encoding property, only a small proportion is active within a motif regardless of its encoding type, indicating motif’s high encoding efficiency. Together, our analysis reveals that motifs and single neurons are parallel encoding schemes at different spatial scales, providing experimental evidence to current population coding theories^(10)^.

## 4 Acknowledgments

We thank Dr. Shoutik Mukherjee for sharing and explaining the published dataset. We thank Dr. Travis Babola for comments on the paper This work was supported by NIH RO1DC017785 (POK)

## 5 Author contributions

L.X, A.C and P.O.K conceived study. M.W performed experiments. L.X analyzed data. A.C and P.O.K supervised the project. L.X, M.W, A.C and P.O.K wrote the paper.

## 7 STAR Methods

### Animal

All protocols and procedures were approved by the Johns Hopkins Institutional Care and Use Committee. In this study, we used 5 male and 3 female adult mice that were the F1 generation of C57BL/6J-Tg(Thy1-GCaMP6s)GP4.3Dkim/J (JAX# 024275) crossed with B6.CAST-Cdh23Ahl+/Kjn mice (JAX#002756), which has minimal hearing loss throughout their lifespan^(70)^. These two mice came from 1 litter, aged between 2 to 3 months when training started, and were less than 6 months old when the imaging data were collected.

### Surgery

All surgeries on our trained animals were performed as previously described^(71)^. We subcutaneously injected 0.1 ml of dexamethasone (2 mg/ml, VetOne) 0.5 to 1 hour before surgery to prevent brain swelling during cranial window implant. We also subcutaneously administered 0.1 ml of dexamethasone (2 mg/ml) right before each surgery. Anesthesia was induced using isoflurane (4% induction, 1.5% maintenance, VetOne). We first removed hair on the top of the head and then removed the skin and soft tissues underneath using disinfected scissors and scalpel blades. Cranial window surgery was then performed on the left side of the skull by first removing muscles on the dorsal and caudal sides to expose the skull. The center of the window location was determined by landmarks on the skull. We then removed a circular area of the skull with a diameter approximately equaling 3.5 mm with a dental drill. We next placed a custom-made cranial window on top after cleaning the exposed brain surface using sterile saline. The window was made with a layer of 3 mm round coverslips (catalog #64–0720-CS-3R, Warner Instruments) stacked at the center of a 4 mm round coverslip (catalog #64–0724-CS4R, Warner Instruments) and secured with optic glue (catalog #NOA71, Norland Products). We then sealed the edge of the window with Kwik-sil (World Precision Instruments) and applied dental cement (C&B Metabond) to secure the window to the skull. A custom-designed headplate was mounted to the skull and dental cement was used to cover the rest of the exposed area of the skull (C&B Metabond). After the surgery, we subcutaneously injected 0.05 ml cefazolin (1 g/vial, West Ward Pharmaceuticals) and 0.03 ml meloxicam (5 mg/ml, VetOne) per gram of mouse body weight, and we placed the mouse under a heat lamp for recovery of 30 min before putting them back in the home-cage. We also subcutaneously administered the same amounts of dexamethasone, cefazolin, and meloxicam on days 1, 2, and 3 post-surgery. We began training 14 days after the surgery. Surgeries on animals of Francis et al. were described elsewhere^(30;29)^

### Behavioral paradigm and training

Mice were water-restricted throughout our training. We would, however, deliver them free water if the body weights were less than 80% of their initial weights at the end of each training day to ensure the survival and normality of the mice. We first put mice into the training chamber for 1 to 3 days for acclimation and would constantly provide water from the spout. The mice were placed inside a plastic tube and were mounted by a custom-designed headplate holder. During acclimation, each mouse would stay in the chamber until it consumed 1 mL of water, which was determined by the change in body weight. The auditory discrimination training would begin after acclimation.

The behavioral paradigm is similar to what was previously described^(53)^. In the two-alternative forced choice tone discrimination task, we trained the mice to turn the wheel to one (e.g. left) direction when hearing a low-frequency (10 kHz) tone and to the other direction (e.g., right) for a high-frequency (40 kHz) tone. All tones were 10 Hz fully amplitude-modulated and were player at 70 dB sound pressure level (SPL) for 1 second. All sound waveforms were generated by UR12 USB Audio Interface (Steinberg) and routed to an ED1 speaker driver (Tucker-Davis Technologies) for presentation using an ES1 speaker (Tucker-Davis Technologies). The speaker was located 10 centimeters away from the animal’s right ear. We calibrated tones of different frequencies in situ with a B&K microphone (Bruel & Kjaer 4944-A). Choices were considered made once the wheel was turned to pass a threshold of 15° in either direction, which would result in a correct or wrong outcome. Trials with no decision made during the choice period (4 seconds after the tone onset) were considered miss trials. Correct trials were rewarded with a drop of water (opening the pump for 200ms, approximately 20 *µ*L). To prevent bias in the wheel-turning direction of mice, we implemented debiasing methods during training once the animals reached around 50% hit rates. We would repeat the trial with the wrong outcome until the animal corrected its behavior. On each training day, the animals must either make 100 correct choices or get trained for one hour, whichever is achieved earlier.

We would start to perform two-photon imaging on the behaving animal once the hit rate reached 80% for three consecutive days. During each behavior session, low-frequency trials and high-frequency trials were presented randomly with a randomized minimum base inter-trial interval (ITI) of 4 to 6 seconds plus time for reward consumption (2 seconds). A new trial would only be delivered when it passed ITI from the end of its previous trial, and the animal held still for at least 1 second (changes in rotary readouts not passing a threshold of 9° in either direction within 1 second). Each behavior session lasted for 33-40 minutes, during which, behavioral results, including rotary readouts, licking activities, timestamps for each trial onset, trial type, and trial outcomes, were continuously recorded.

The behavior paradigm and training procedure of the published dataset are described in^(29)^. In short, mice were trained to discriminate different sounds through licking a water spout. Each trial began with 1 s of silence, followed by a 55 dB SPL amplitude modulated (8 Hz) tone lasting for 1 s. The total length of each trial was 4.5s. The mice were trained to lick the water spout after the onset of a low-frequency sound (7 kHz or 9.9 kHz) and to avoid licking after the onset of a high-frequency sound (14 kHz or 19.8 kHz). The tone frequency was randomized across trials. Trials were separated by a random 5–9 s inter-trial interval (ITI). Each trial’s behavioral response was categorized as a hit (licking after a low-frequency tone), miss (no licking after a low-frequency tone), false alarm (licking after a high-frequency tone), or correct rejection (no licking in response to a high-frequency tone). Incorrect behavioral responses were punished with an 8 s time-out added to the ITI. To reduce impulsive licking and improve task performance, the mice were trained to delay behavioral responses until 0.5 seconds after the onset of a target tone to be rewarded with a water droplet. Mice were trained on the task until hit rates were consistently above 70% and then imaged during behavior. Mouse health was monitored daily by a skin turgor test and checking that body weight remained above 80% of the initial off-study weight.

### Imaging

On our trained animals, two-photon imaging was performed using a Bruker Ultima 2Pplus microscope with a 16 × objective (Nikon, 0.80 NA, 3.0 mm WD). The laser (Coherent Discovery) was tuned at 920nm to excite the green fluorescence of the calcium (Ca^2+^) indicators^(72)^. In each imaging session, we recorded the Ca^2+^ activity from the primary auditory cortex (A1) at a specific depth between 100*µ*m and 400*µ*m below the brain surface S5. A1 was identified by the axis of the tonotopy gradient based on previously described methods using widefield imaging^(56)^. Each imaging field of view was captured by resonant galvo scanning of a 512 × 512 pixel-sized area at 30 Hz with a pixel size of 1.084*µ*m. The onset of each trial was signaled by a trial trigger sent through the GPIO extension board of the Raspberry Pi computer that functioned as the delivery of behavioral trials. The trial triggers were recorded through the PrairieView software (version 5.8) together with the imaging frame triggers as voltage recording signals and were used to synchronize the imaging data and behavioral data.

In the published dataset, mice were implanted with a 3 mm chronic window over the auditory cortex. During each trial, two-photon(2P) calcium imaging was recorded in a selected window(370 × 370 *µm*) in primary auditory cortex (A1) for each mouse. Imaging frames of 512 × 512 pixels (pixel size 0.72 *µm*) were acquired at 30 Hz by bidirectional scanning of a scanning microscope (Bergamo II series, B248, Thorlabs). For each imaging session, neurons were located and the cell and neuropil fluorescence activities were estimated for each neuron using prior methods. For details see^(29;30)^.

### Data Preprocessing

For each of our imaging sessions, we ran Suite2P (https://suite2p.readthedocs.io/en/latest/) to automatically identify cells and then used SEUDO^(73)^ to remove erroneous cells or artifacts with manual inspection. We did not correct any cells found by Francis et al. since they were labeled manually. Once we had the cell profiles and their fluorescence traces, we took a few pre-processing steps to make the data fit into the motif-detection algorithm. We first corrected the neuropil fluorescence activities with 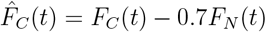, where *F*_*C*_(*t*) were the uncorrected individual neuron fluorescence traces and *F*_*N*_ (*t*) were the neuropil fluorescence traces around that cell. We then found the baseline fluorescence activity *F* as the 25th percentile of the 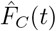 for each trial and calculated the Δ*F/F* for each neuron. To account for heterogeneous noise levels, the Δ*F/F* trace of each neuron was re-scaled to have unit robust standard deviation^(74)^. We estimated the peak signal-to-noise ratio(PSNR) of each neuron as the square of the peak signal(95th percentile of Δ*F/F*) over the variance of Δ*F/F*. Since both the SeqNMF algorithm and our adapted algorithm require the input data matrix to be non-negative and sparse(i.e., most entries are zero), we only preserved the true calcium transients(continuous Δ*F/F* activities above 2 and lasting for at least 0.2s) in the data and set all the other signals to zero (Fig S2A,B).

To improve the efficiency of the motif-detection algorithms, we defined good neurons as neurons whose PSNR were greater than 3 and have at least 8 instances of calcium transients in the data(i.e., 4 times in the training data and 4 times in the test data). the PSNR threshold was determined based on the distribution of each neurons’s PSNR(Fig S2C). We only keep the good neurons in each session for motif analysis. To ensure the algorithm could find meaningful motifs from enough good neurons and different trial types, we restricted our analysis to the sessions that have at least 10 good neurons and at least 10 trials in each sound frequency(Table S1).

Note that the Split Bregman Iteration (SBI) updating rule(4) is sensitive to the scaling of the data. To make the parameter selection more efficient and to guarantee fair comparison between different algorithms, we normalized the training data to have unit Frobenius norm 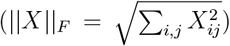 before running any motif detection algorithm.

### Cosine similarity between neural population activity

Given the population activity in two different trials *X*_*i*_, *X*_*j*_ ∈ ℝ^*N* ×*T*^, we vectorized the matrices and computed the cosine similarity between them: 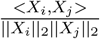

### SeqNMF algorithm

Given a data matrix **X** ∈ ℝ^*N* ×*T*^, SeqNMF tries to decompose it into the sum of convolution of *K* different motifs (factors) **W** ∈ ℝ^*N* ×*K*×*L*^ and their temporal loadings **H** ∈ ℝ^*K*×*T*^ (Fig. 2A):

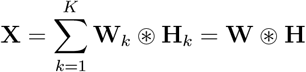

where *N* is the number of neurons, *K* is the number of different motifs, *L* is the length of each motif. For real data that contains noise, the algorithm seeks to find the optimal factorization 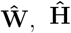 by solving:

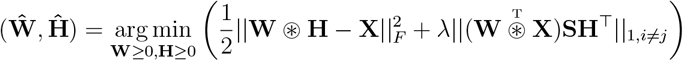

where the first term is the reconstruction error and the second term is the regularization of the motifs to be non-redundant. *λ* represents a trade-off parameter between the two terms.

Note the cost function is only convex in **W** and **H** separately but not jointly, SeqNMF alternate solves over **W** and **H** (2,3) through a multiplicative updating rule(MUR):

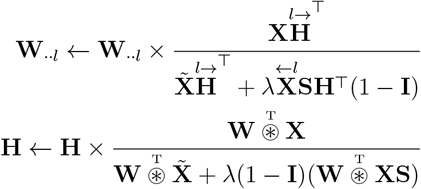

MUR is a special case of gradient descent updates and will end up reaching a local minimum of the optimization program. For detailed derivation see^(37;46;32)^.

### Split Bregman Iteration

We applied Split Bregman Iteration (SBI)^(45;44)^ to solve (2),(3) in the SeqNMF program. Split Bregman methods have been proposed as a powerful tool to solve a very broad class of L1-regularized optimization problems. Particularly, SBI has been successfully applied to solve the analysis based sparsity prior problem:

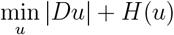

where *D* is linear operator on *u* and *H*(·) is a smooth convex function, such as 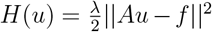. Note that the first term of (2) is a smooth, convex function of **H**, the second term of (2) can be written as *D***H** where *D* is a linear operator on **H** (element-wise) and |·| denotes the l1 norm. By applying the SBI approach in^(44)^ in solving (2), we get (4). Similarly, we can derive the SBI method for solving (3):

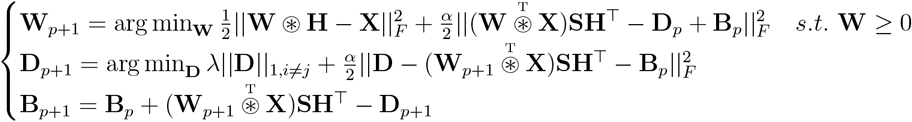

The first line is a convex optimization program with non-negative constraints. The second line is a LASSO step. The programs were solved using TFOCS^(75)^ in MATLAB. In practice, we found that the first line converged slowly when using the default solver in TFOCS (Auslender and Teboulle’s method^(76)^). We thus changed the solver to N83 (^(77;78;79)^) and applied the restart method(^(80)^) to accelerate. We set the restart step size to 50 and the maximum iterations of the solvers to 500 to achieve rapid convergence. We also observed that the iteration (4) converged fast so we a set a maximum repeats of (4) to 10. The iteration could end in advance when the difference of the optimization variable **W** or **H** between iterations(measured as the frobenius norm) is below tolerance. For a regular session, it takes about 15 minutes to run SBI and find motifs on a desktop (Ryzen 5900 12-core, 64G memory).

### Comparing MUR and SBI on Simulated data

To compare the performance of the original SeqNMF algorithm based on MUR and our devised algorithm based on SBI, we simulated a dataset that consisted of 200 trials. 100 trials were put to the training set and 100 trials were put to the test set. 10 different motifs randomly occur 1-10 times in both the training and test set. Each motif consists of 5 neurons that consecutively activate with a positive transient activity. To simulate calcium transients, we set the transient to be the convolution of a square wave and an exponential decay function. We applied various noise types to the data, including additive spike noise, probabilistic participation, temporal jitter, temporal warping, and shared neurons between motifs. The first four types were the same as^(37)^. For shared neurons between motifs, we set 3 neurons to be shared between two different motifs(See Figure 2).

For a fair comparison, we tuned the hyperparameter *λ* to be optimal for the original SeqNMF algorithm using the same procedure described in^(37)^. We then used the same *λ* in MUR and manually tuned *α*_**H**_ and *α*_**W**_ to some sub-optimal values. For the simulation in Figure 3, *λ* was set to 0.1(Fig S6), *α*_**H**_ was set to 1e-3 and *α*_**H**_ was set to 1e-6. The complete hyperparameter setups under other noise conditions are provided in Table S2.

### Significance test of motifs

To test if the motifs we extracted were significantly present in the test data, we followed a procedure similar to^(37)^. In brief, we measured the overlaps of each motif with the test data as 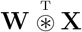. The overlap will be high when the motif occurs, forming a long-tail distribution across all the time bins (Fig S5A). To ensure that we were extracting meaningful motifs with stereotypical spatiotemporal Xstructure, we constructed null motifs by circularly shifting the time course of each neuron by a random amount between 0 to L. We could then get a distribution of 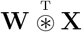 for each null motif we built (Fig S5B). We realized that when the motif appeared very few times in the data and consisted of very few neurons, there were too few time points when 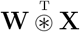 reached a large value. Thus, the skewness statistic used in the SeqNMF paper was not sensitive enough to capture the difference in the distribution of 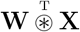 between the true motif and null motifs. We thus focused on the ‘tail’ of the distribution (i.e., the top 5% largest values of the distribution) and tested whether the mean of the tail of the true motif would be greater than the chance level(i.e., mean of the tail of the null motif, Fig S5C).

### Parameter Selection in Split Bregman Iteration

Three hyperparameters *λ, α*_**H**_ and *α*_**W**_ are involved in the SBI algorithm. First, we noticed that the motifs themselves did not need to be sparse, so we set *α*_**W**_ to a small value (1e-6). Increasing *α*_**H**_ or *λ* will increase the weight of the regulariation term, driving the algorithm to produce sparser solutions. We designed a procedure based on the idea that the algorithm should produce motifs that have precise timing (sparse enough in **H**) and at the same time capture as many stereotypical spatiotemporal structures(motifs) as in the data. To evaluate the sparseness of **H**, we denoted the proportion of non-zero entries in 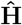 as *S*. To evaluate the proportion of spatiotemporal structures in the data captured by the algorithm, we decomposed the data matrix into two parts:

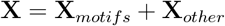

where 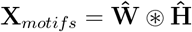 is the ‘motifs’ part of the data reconstructed by the algorithm and **X**_*other*_ is the remaining part of the data that is not stereotyped(Fig S7A). To ensure **X**_*other*_ does not contain any stereotypical spatiotemporal structures, we computed the peak of the cross-correlation between each pair of neurons’ traces within a certain time window in both **X** and **X**_*other*_, resulting in two cross-correlation matrices *C*_**X**_ and *C*_*Other*_. The off-diagonal part of *C*_**X**_ formed a long-tail distribution as only a small fraction of neurons were involved in motifs in real data(Fig S7B). After subtracting the ‘motif’ part of the data **X**_*motifs*_, the large values of the off-diagonal part of *C*_*Other*_ should vanish. We thus used

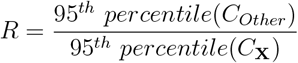

to quantify the proportion of ‘motif’ structure that was not captured by the algorithm (Fig S7B). When we plotted *S* and *R* as a function of *α*_**H**_ and *λ*, we saw decrease in *S* and increase in *R* as we increase *α*_**H**_ or *λ*(Fig S7C). In between there was a point when both *S* and *R* were low, indicating a good capture of motifs with temporal precision (annotated with bold black box). We also plotted the reconstruction and regularization costs as a function of *α*_**H**_ and *λ* as in the SeqNMF paper. The optimal hyperparameter point we found using our procedure was consistent with the optimal point of the SeqNMF paper where reconstruction and regularization costs were balanced(Fig S7C).

### Determine the activation time of motifs

After we identified the temporal loading of each motif **H**_*k*_ with the algorithm, we applied median filtering with a window of 5 time bins and extracted all the non-zero segments. For each segment, we used its geometric mean as the exact motif activation time. We defined the magnitude of each occurrence as the integral of its corresponding segment. The occurrences whose magnitudes were too small (less than 5% of the norm of **H**_*k*_) were dropped.

### Determine active neurons in motifs

The detected motifs can recruit neurons that participate in the motif with probability or neurons with high spontaneous activities. To distinguish neurons that are truly active in the motif, we developed the following significance test for each neuron. Given the exact activation time *t*_1_, *t*_2_, …, *t*_*R*_ of a certain motif, we randomly sampled the same amount of time points as ‘null activation time’. We evaluated the overlaps between motif *W*_*k*_ with the data segments at true activation times **X**_*t*1_, **X**_*t*2_, …, **X**_*tR*_ (element-wise multiplication), where each segment has the same dimension as **W**_*k*_ ∈ ℝ^*N* ×*L*^. We summed up the overlap along the time dimension, resulting in a distribution of overlaps for each neurons. Similarly, we can compute the distribution of overlap for each ‘null activation time’, resulting in a ‘null overlap’ for each neuron. We performed a Wilcoxon rank sum test between the distribution of true overlap and all the ‘null overlaps’ for each neuron to determine if the neuron is significantly active in the motif.

### Spatial distribution of motif active neurons

To test if the motif active neurons within motifs are spatially clustered or regularly ordered, we used Clark-Evans R index^(48)^ to measure the clustering of active neurons within field of view. The Clark-Evans Aggregation Index provides a ratio R between the observed average nearest neighbor distance (NND) and expected average NND under complete spatial randomness (CSR). *R <* 1 means neurons are spatially clustered, *R >* 1 means neurons are regularly ordered, and *R* ≈ 1 means neurons are randomly distributed. Since the number of active neurons is small, we used bootstrap to simulate the distribution of R under null hypothesis (*R* = 1) and tested if the R indices of active neurons are significantly deviated from 1. To ensure statistical power, we only tested on motifs with at least 5 active neurons. Bonferroni correction was conducted to account for multiple tests on multiple motifs. Among all the motifs in both datasets, we found no evidence for any motif with neurons with clustered or regularly spaced active neurons (Fig. S11). However, given the number of active neurons is small, failure to reject CSR should be interpreted cautiously.

### Determine the onset time within motif

The activation time of a motif may not give us the precise onset time of the motif, since the motif last for *L* time bins and active neurons may not be active at the very beginning of the motif. Thus, we define the onset time of the motif as the earliest onset time of all the active neurons. To determine the onset time of each active neurons, we normalized each row of **W** to unit norm and find the first point above 0.05 in each row. We plot all the significant motifs and their onsets to justify our criteria(Fig S8). By comparing the motif activation time, the onset time within each motif against trial onsets, we could determine the precise motif onsets within each trial (Fig 4C).

### Quantify the percent variance explained by motifs and neurons

We adopted a similar procedure as SeqNMF to quantify the percent variance explained by each motif. Given a motif **W**_*k*_ and its temporal loading **H**_*k*_, the total variance of the data **X** is ∑ **X**^2^, the percent of total variance unexplained by this motif is (**X** − **W**_*k*_ ⊛ **H**_*k*_)^2^. Thus, the percent of variance explained by the motif is:

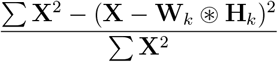

Similarly, within one motif **W**_*k*_ ∈ ℝ ^*N* ×*L*^, the variance of the *n*^*th*^ neuron is *V*_*n*_ = ∑ _*l*_ **W**_*k*(*n,l*)_. The percent of variance explained by the *n*^*th*^ neuron is 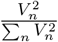

### Computing motif encoding indices of task variables

To quantify a motif’s selectivity to a certain task variable(stimulus, choice, licking, stimulus & choice), we built an encoding index for the variable based on the motif’s occurrence frequencies in all trial types. Take Francis et al. dataset as an example, we first computed a 4 × 2 × 2 probability distribution matrix *P* (*S, C, M*) from all the trials in that session. The first dimension indicates the trial’s stimulus (one of the four frequencies), the second dimension indicates the animal’s choice in that trial (C=1 means correct, C=2 means error), the third dimension indicates whether the motif occurred (M=1) or not (M=2) in that trial. For example, *P* (*S* = 2, *C* = 1, *M* = 1) is the proportion of trials in which f2 was played, the mouse made correct choices and the motif occurred. For simplicity, we denote *P* (*S* = *i, C* = *j, M* = *k*) as *P* (*S*_*i*_, *C*_*j*_, *M*_*k*_). We then computed the conditional probability of the motif occurring in trials with a certain stimulus being played *P* (*M*_1_ |*S*_*i*_), *i* = 1, 2, 3, 4 and sorted them in descending order. We defined the stimulus-encoding index of the motif as the largest conditional probability minus the second-largest conditional probability, which ranges between 0 to 1. A high encoding index(close to 1) indicates the motif exhibits strong sound selectivity, while a low encoding index(close to 0) indicates weak selectivity. Similarly, we could also measure the conditional probability of the motif occurring in trials with a certain choice *P* (*M*_1_|*C*_*j*_) and the conditional probability of the motif occurring in trials with a stimulus and choice jointly *P* (*M*_1_| *S*_*i*_, *C*_*j*_), getting the choice encoding and joint-encoding indices respectively. For lick-encoding index, we simply change the choice dimension to licking and built the probability distribution matrix *P* (*S, L, M*) in a similar way(L=1 means licking, L=2 means not licking, which takes the same value as C in low-frequency trials but the opposite value in high-frequency trials).

### Determine movement onsets and offsets in wheel-turning trials

To determine the onsets and offsets of the wheel movement in each trial, we defined the ‘choice direction’ of a certain trial as the direction with the maximal absolute turning speed within that trial. As the animal could make spontaneous movements, we defined the onset of wheel movement as the time when the animal last turned into the ‘choice direction’ after sound onset and before tuning with the maximal speed, and defined the offset of wheel movement as the first time when the animal stopped turning into the ‘choice direction’ (or start turning to the reverse direction) after turning with the maximal speed(Fig 5C).

### Test motif encoding of task variables

Given the animals were well trained for the sound discrimination tasks, the number of miss and error trials is limited, Additionally, the motif’s encoding of sound is correlated with the encoding of choice, posing challenges to the test of motif encoding of different task variables. To address the challenge, we applied a series of Fisher’s exact tests (https://www.mathworks.com/help/stats/fishertest.html),

which has been shown to work well on small samples. To disentangle stimulus and choice encoding, we first created a 2-by-2 contingency table based on stimulus types and motif activations (regardless the choice being made) to test if the motif significantly encoded a certain stimulus. We then tested the encoding of a certain choice for each motif by building another contingency table based on all the trials with one frequency. The motif is deemed as joint encoding if it significantly encodes choice *C*_*j*_ conditioned on stimulus *S*_*i*_, but not conditioned on other stimuli. It is deemed as choice-encoding (or lick-encoding) only if it significantly encodes choice *C*_*j*_ conditioned on all stimuli. Bonferroni correction was conducted to account for multiple tests on multiple motifs and variables. Given the small number of trials in some trial types, it is likely that sometimes the minimum p-value Fisher’s exact test can produce is larger than the significance level *α*. In this case, the motif’s encoding of a certain variable is untestable, which is marked as X in table S5 and table S4.

To test the encoding of a specific event variable within trial (e.g., reward, movement onset, movement offset), we tested if the motif onsets are better aligned to the events than to the sound onsets among the trials the event occurred using two-sample F-test (https://www.mathworks.com/help/stats/vartest2.

### Compare motif neurons against GC-linked neurons

Francis et al. performed GC analysis on 20 intersection information (II) neurons in 12 out of 34 sessions. Intersection information(II) measures the part of the sensory information that is used to inform behavioral readout^(51)^. Detailed procedure on computing II and and their significance are described in^(29)^. Granger causality(GC) analysis measures the predictive influence of the past activity of one neuron on the present activity of another. Details on inferring the GC networks are described in^(29)^ and ^(30)^. For comparison, they also inferred GC networks on the 20 best responding neurons (Fig S8B). All the GC networks are inferred on Hit, Miss, Correct Rejection(CR) and False Alarm(FA) trials separately. Since motifs were identified on all the trials instead of different types of trials, we grouped each task-encoding motif into one of the trial types in which it has the maximum conditional probability of occurrence (measured in a similar way as in computing encoding indices). We then computed the dice coefficients between the GC-linked neurons and motif active neurons. Note that the GC-network were estimated at two time scales(i.e., the duration of recent history in which interactions are quantified, referred as the ‘integration window’), short(223ms) and long(1033ms). Given our motifs span 100 time bins (3.33s), we made the comparison for both time scales. The GC links can have positive and negative values, representing facilitative or suppressive interactions respectively. Given that the neural activities within motifs are all positive, we compared the motif neurons against positive GC-linked neurons(top rows of Fig 4E and Fig S8B) as well as all GC-linked neurons (bottom rows of Fig 4E and Fig S8B).

### Balancing index between different trial types and its effect on joint-encoding motif count

We applied three balance metrics to quantify the distribution of three trial types conditioned on a certain stimulus.

#### Entropy

*Balance* = − ∑_*i*∈{corr,miss,error}_ *p*_*i*_ log_3_ *p*_*i*_

#### Max difference from idea

*skew* = max(|*p*_*corr*_ − 1*/*3|, |*p*_*miss*_ − 1*/*3|, |*p*_*error*_ − 1*/*3|)

#### Gini Coefficient

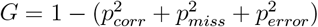

To explore if the balance between different trials have a significant effect on the identification of joint-encoding motifs, we applied two-sample one-sided t-tests for each of these metrics between sessions with and without joint-encoding motifs. We observed a significant effect of trial balance within the low frequency trials (p=6.16E-5 for entropy, p=2.39E-6 for skew, p=1.09E-5 for Gini Coefficient) and a weak effect within the high frequency trials (p=0.03 for entropy, p=0.27 for skew, p=0.26 for Gini Coefficient). Overall our analysis show that even distribution of trial types contribute to the identification of joint-encoding motifs.

### Single neuron encoding analysis

#### Stimulus-encoding neurons

Neurons in A1 can respond to sound onsets or offsets ^(56)^. To identify stimulus-encoding neurons, we compared the neural activities (mean dF/F) in three sound-related time windows (0-0.5 second after sound, onset, 0.5-1 second after onset, and 0-0.5 second after offset) against baseline (0-1 second before sound onset, i.e., the first second in each trial) using paired sample t-test (https://www.mathworks.com/help/stats/ttest.html) at Bonferroni corrected significance level 0.05/N. A neurons is deemed as stimulus-encoding if it shows significantly larger dF/F in any of the three windows. The stimulus encoded by the neuron is defined as the stimulus under which it has largest mean responses.

#### Choice-encoding and joint-encoding neurons

We observed that, in most trials, the decisions were made during 0-1.5 seconds after sound onsets (Fig. S12B). Considering the temporal delay of calcium signal, we thus used the difference between mean dF/F over 0-2 seconds after sound onset and mean dF/F over 0-1 second before sound (baseline) as the neural response in each trial. Neural responses can be driven by stimulus or choice signal. To disentangle their effects, we took a similar approach as in motif-encoding analysis to identify choice-encoding and joint-encoding neurons. Specifically, we grouped the trials under different stimuli (low and high frequency) separately and tested if the choice had a significant effect on the neuron’s response conditioned on each stimulus. Since very small samples of certain trial types (miss or error) can result in low statistical power, we only tested the effect of choices with at least 5 trials conditioned on a certain stimulus. If all the three choices (Hit, Miss, Error) were testable, we applied one-way ANOVA (using Satterthwaite method to compute the degree of freedom) to test the effect of choice on neural response. If only two choices were testable, we applied two-sample t-test to compare the neural responses between trials with the two choices. Both tests were conducted at Bonferroni corrected significance level 0.05/N. The choice encoded by the neuron (conditioned on a certain stimulus) is defined as the choice under which it shows the greatest mean neural response. To ensure fair comparison against motif encoding (which only focused on positive neural responses), we only kept neurons whose largest responses over all testable choices were greater than 0 (one-sample t-test). Similar to what we have done for motif-encoding, a neuron is deemed choice-encoding only if it encodes the same choice under both frequencies. It is deemed as joint-encoding of stimulus and choice if it only encodes one choice under one frequency or encodes different choices under two frequencies. All the choice-encoding and joint-encoding neurons were excluded from the stimulus-encoding category (See Fig. S12C for examples).

#### Reward-encoding neurons

We adopted a similar approach to identify reward-encoding neurons by testing whether a neuron’s calcium transients were time-aligned to the reward onsets. Since reward-encoding may also happen jointly with the presence of certain sound, we tested if the neuron’s true transients given by SEUDO were time-aligned to the reward timing conditioned on two frequencies separately. Among all the session, we did not observe any reward-encoding neurons, even conditioned on a certain stimulus.

#### Movement-encoding neurons

To disentangle wheel movement from stimulus response and decision making, we tested if the wheel movement during inter-trial intervals (0-3 seconds before trial onsets) were correlated with neural activity dF/F. Specifically, we computed the cross correlation between the smoothed neuron’s dF/F and the smoothed movement speed (window length = 5) during each inter-trial intervals. The maximum time lag for cross-correlation is set to 15 time bins. The movement speed can be signed (distinguishing left and right movement) or absolute movement speed. By looking into the distribution of these cross correlations and their peak times, most neurons show weak correlations and the peak of correlation usually occurs before 0 (Fig S12), indicating weak response to motion. To test the significance of these correlations, we randomly shuffled the trial ids to create numerous ‘null speed trajectories’ during ITI, and computed the cross correlations between neural activities and these ‘null speed trajectories’. We then measured if the correlation at each time lag is significant at the Bonferroni-corrected significance level.

### Comparing motif encoding against single neuron encoding properties

To investigate if the active neurons within motifs share the same encoding properties as the motifs, we compute the proportion of overlap between motif active neurons with all encoding neurons. Specifically, denote the set of active neurons of a motif with encoding property i as *M*_*i*_, denote the set of all neurons with encoding property j as *N*_*j*_, we compute two ratios: the overlap over all motif neurons 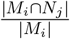 (the mean of this ratio over all motif is plotted in Fig. 6D top) and the overlap over all encoding neurons 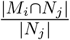 (the mean of this ratio over all motif is plotted in Fig. 6D bottom). Similarly, for the sets of neurons from two different motifs with encoding properties i and j, we can compute their overlap as 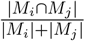 in Fig. 6E.

## Supplement Materials

**Table S1.**
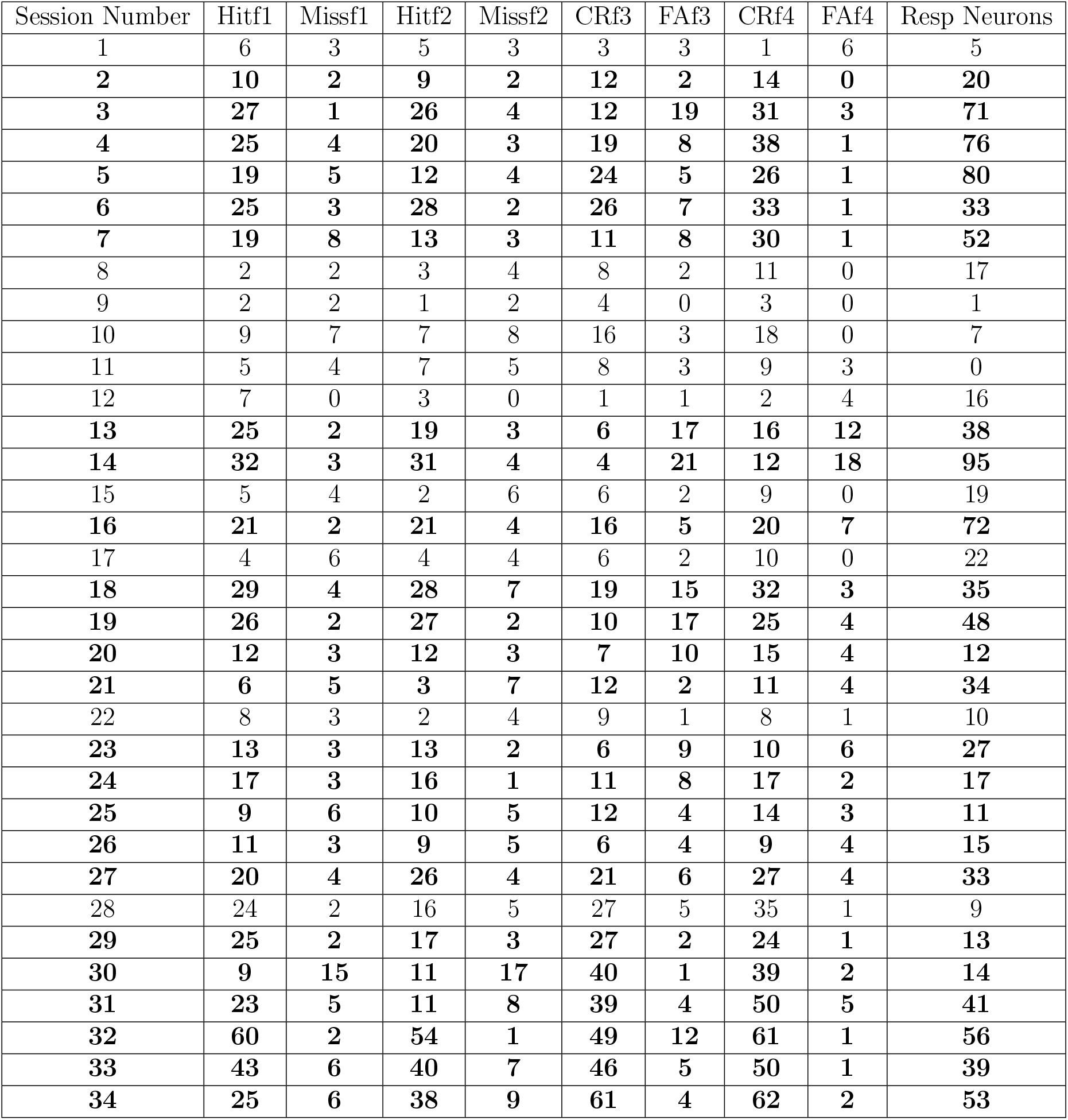
Trials and neurons statistics of Francis et al. two-photon dataset Number of different trials and responsive neurons in all the sessions in Francis et al. dataset. Bold sessions were selected for motif analysis

| Session Number | Hitf1 | Missf1 | Hitf2 | Missf2 | CRf3 | FAf3 | CRf4 | FAf4 | Resp Neurons |
| --- | --- | --- | --- | --- | --- | --- | --- | --- | --- |
| 1 | 6 | 3 | 5 | 3 | 3 | 3 | 1 | 6 | 5 |
| **2** | **10** | **2** | **9** | **2** | **12** | **2** | **14** | **0** | **20** |
| **3** | **27** | **1** | **26** | **4** | **12** | **19** | **31** | **3** | **71** |
| **4** | **25** | **4** | **20** | **3** | **19** | **8** | **38** | **1** | **76** |
| **5** | **19** | **5** | **12** | **4** | **24** | **5** | **26** | **1** | **80** |
| **6** | **25** | **3** | **28** | **2** | **26** | **7** | **33** | **1** | **33** |
| **7** | **19** | **8** | **13** | **3** | **11** | **8** | **30** | **1** | **52** |
| 8 | 2 | 2 | 3 | 4 | 8 | 2 | 11 | 0 | 17 |
| 9 | 2 | 2 | 1 | 2 | 4 | 0 | 3 | 0 | 1 |
| 10 | 9 | 7 | 7 | 8 | 16 | 3 | 18 | 0 | 7 |
| 11 | 5 | 4 | 7 | 5 | 8 | 3 | 9 | 3 | 0 |
| 12 | 7 | 0 | 3 | 0 | 1 | 1 | 2 | 4 | 16 |
| **13** | **25** | **2** | **19** | **3** | **6** | **17** | **16** | **12** | **38** |
| **14** | **32** | **3** | **31** | **4** | **4** | **21** | **12** | **18** | **95** |
| 15 | 5 | 4 | 2 | 6 | 6 | 2 | 9 | 0 | 19 |
| **16** | **21** | **2** | **21** | **4** | **16** | **5** | **20** | **7** | **72** |
| 17 | 4 | 6 | 4 | 4 | 6 | 2 | 10 | 0 | 22 |
| **18** | **29** | **4** | **28** | **7** | **19** | **15** | **32** | **3** | **35** |
| **19** | **26** | **2** | **27** | **2** | **10** | **17** | **25** | **4** | **48** |
| **20** | **12** | **3** | **12** | **3** | **7** | **10** | **15** | **4** | **12** |
| **21** | **6** | **5** | **3** | **7** | **12** | **2** | **11** | **4** | **34** |
| 22 | 8 | 3 | 2 | 4 | 9 | 1 | 8 | 1 | 10 |
| **23** | **13** | **3** | **13** | **2** | **6** | **9** | **10** | **6** | **27** |
| **24** | **17** | **3** | **16** | **1** | **11** | **8** | **17** | **2** | **17** |
| **25** | **9** | **6** | **10** | **5** | **12** | **4** | **14** | **3** | **11** |
| **26** | **11** | **3** | **9** | **5** | **6** | **4** | **9** | **4** | **15** |
| **27** | **20** | **4** | **26** | **4** | **21** | **6** | **27** | **4** | **33** |
| 28 | 24 | 2 | 16 | 5 | 27 | 5 | 35 | 1 | 9 |
| **29** | **25** | **2** | **17** | **3** | **27** | **2** | **24** | **1** | **13** |
| **30** | **9** | **15** | **11** | **17** | **40** | **1** | **39** | **2** | **14** |
| **31** | **23** | **5** | **11** | **8** | **39** | **4** | **50** | **5** | **41** |
| **32** | **60** | **2** | **54** | **1** | **49** | **12** | **61** | **1** | **56** |
| **33** | **43** | **6** | **40** | **7** | **46** | **5** | **50** | **1** | **39** |
| **34** | **25** | **6** | **38** | **9** | **61** | **4** | **62** | **2** | **53** |

**Table S2.** Optimal hyperparameters of SBI for simulated data with different noise type

| Noise type | $\lambda$ | $\alpha_W$ | $\alpha_H$ |
| --- | --- | --- | --- |
| Clean | 0.005 | 1e-06 | 1e-02 |
| Additional Noise | 0.003 | 1e-05 | 1e-02 |
| Probabilistic participation | 0.01 | 1e-06 | 1e-03 |
| Temporal Jittering | 0.01 | 1e-06 | 1e-03 |
| Temporal Warping | 0.01 | 1e-06 | 1e-03 |
| Shared Neurons | 0.01 | 1e-06 | 1e-03 |

**Table S3.**
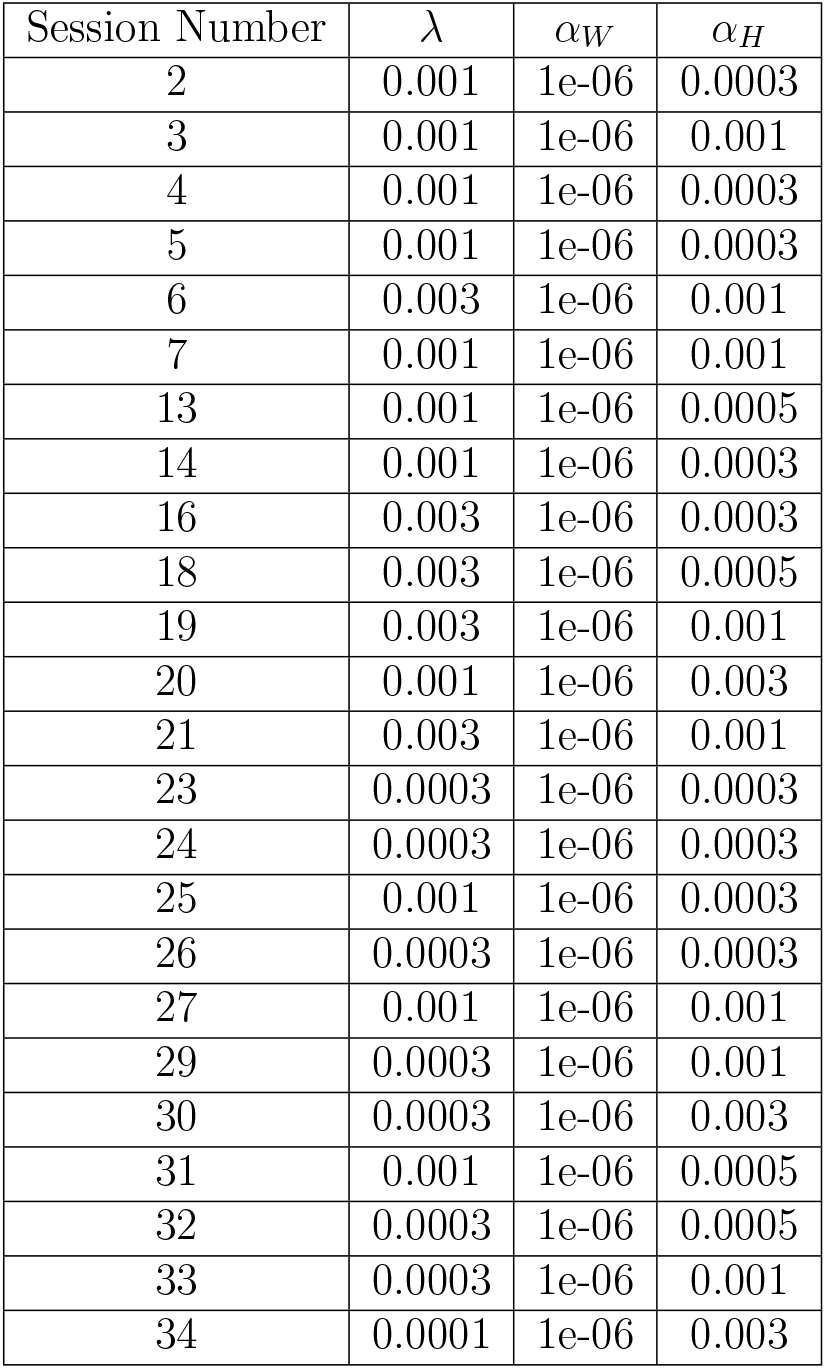
Optimal hyperparameters of SBI for each session

**Table S4.** Motif encoding of different task variables in Francis et al. dataset

| Session | Stimulus | Choice | Licking | Reward | Joint |
| --- | --- | --- | --- | --- | --- |
| 2 | #4,10 | X | X | N | N |
| 3 | #4 | X | X | N | N |
| 4 | N | X | X | N | N |
| 5 | #1,2,4,5 | X | X | N | #9 |
| 6 | N | X | X | N | N |
| 7 | #1,3,6 | X | X | N | N |
| 13 | #1,2 | X | X | N | N |
| 14 | #1,2,3,8 | N | N | N | N |
| 16 | #1 | X | X | N | N |
| 18 | N | X | X | N | N |
| 19 | #1,4 | X | X | N | N |
| 20 | N | X | X | N | N |
| 21 | N | X | X | N | N |
| 23 | #1 | X | X | N | N |
| 24 | #1,3 | X | X | N | N |
| 25 | N | N | N | N | #1 |
| 26 | #1 | N | N | N | N |
| 27 | N | N | N | N | N |
| 29 | N | N | N | N | N |
| 30 | N | N | N | N | N |
| 31 | #2 | N | N | N | N |
| 32 | #1,2,6 | X | X | N | #4 |
| 33 | N | X | X | N | N |
| 34 | #3 | N | N | N | N |

**Table S5.** Motif encoding of different task variables in 2AFC sound discrimination task

| Mouse | Session | Depth($\mu\text{m}$) | FOV | Stimulus | Choice | Joint | Movement+ | Movement- | Reward |
| --- | --- | --- | --- | --- | --- | --- | --- | --- | --- |
| 1M | 1 | 220 | A1 | #1,2,3 | N | N | N | N | N |
| 1M | 2 | 330 | A1 | #1,2,4,8 | N | N | N | N | N |
| 1M | 3 | 290 | A1 | #4,6,7 | X | N | N | N | N |
| 1M | 4 | 290 | A1 | N | N | N | N | N | N |
| 2F | 5 | 220 | A1 | #1,5 | X | N | N | N | N |
| 2F | 6 | 330 | A1 | N | N | #1 | N | N | #4,6 |
| 2F | 7 | 160 | A1 | #3,4 | N | #1 | N | N | N |
| 3M | 8 | 310 | DM | #2,3,9 | N | #1,6 | N | N | N |
| 3M | 9 | 210 | DM | #6,7 | N | #1 | N | N | N |
| 3M | 10 | 170 | DM | #1 | #5 | N | N | N | N |
| 3M | 11 | 350 | DM | #2,4 | N | #3,8 | N | N | N |
| 3M | 12 | 340 | DM | #1,2,6 | N | N | N | N | N |
| 4M | 13 | 235 | DM | N | N | N | N | N | N |
| 4M | 14 | 100 | DM | #2,3 | N | N | N | N | N |
| 4M | 15 | 200 | DM | #1,2,5 | N | #1 | N | N | N |
| 4M | 16 | 340 | DM | #1 | N | N | N | N | N |
| 4M | 17 | 150 | DM | N | #1 | #2,3 | N | N | N |
| 4M | 18 | 275 | DM | N | N | N | N | N | N |
| 5F | 19 | 160 | DM | #1,2 | N | N | N | N | N |
| 6M | 20 | 175 | DM | #1,6 | N | N | N | N | N |
| 6M | 21 | 330 | A1 | #1,4 | X | N | N | N | N |
| 6M | 22 | 280 | DM | #1,2,3,4 | X | N | N | N | N |
| 6M | 23 | 210 | A1 | #1 | X | N | N | N | N |
| 6M | 24 | 365 | DM | #2,5 | N | N | N | N | N |
| 6M | 25 | 275 | A1 | #1 | X | N | N | N | N |
| 7M | 26 | 265 | A1 | N | X | N | N | N | N |
| 7M | 27 | 150 | A1 | #1,3,5,6 | N | N | N | N | N |
| 7M | 28 | 210 | DM | #2,5,6,7,8 | X | N | N | N | N |
| 7M | 29 | 355 | A1 | N | N | #1,3 | N | N | N |
| 7M | 30 | 310 | DM | #1,7,8 | X | #2 | N | N | N |
| 7M | 31 | 180 | A1 | #3,4,5 | N | N | N | N | N |
| 7M | 32 | 255 | DM | #10 | N | N | N | N | N |
| 7M | 33 | 305 | A1 | #9 | N | #1 | N | N | N |
| 7M | 34 | 355 | A1 | N | N | N | N | N | N |
| 7M | 35 | 165 | A1 | #1 | N | #2 | N | N | N |
| 7M | 36 | 200 | A1 | #9 | N | #1 | N | N | N |
| 8F | 37 | 195 | DM | #1 | N | N | N | N | N |
| 8F | 38 | 340 | DM | #3 | N | N | N | N | N |
| 8F | 39 | 285 | DM | N | N | N | N | N | N |
| 8F | 40 | 235 | DM | #4 | N | N | N | N | N |

**Figure S1.**
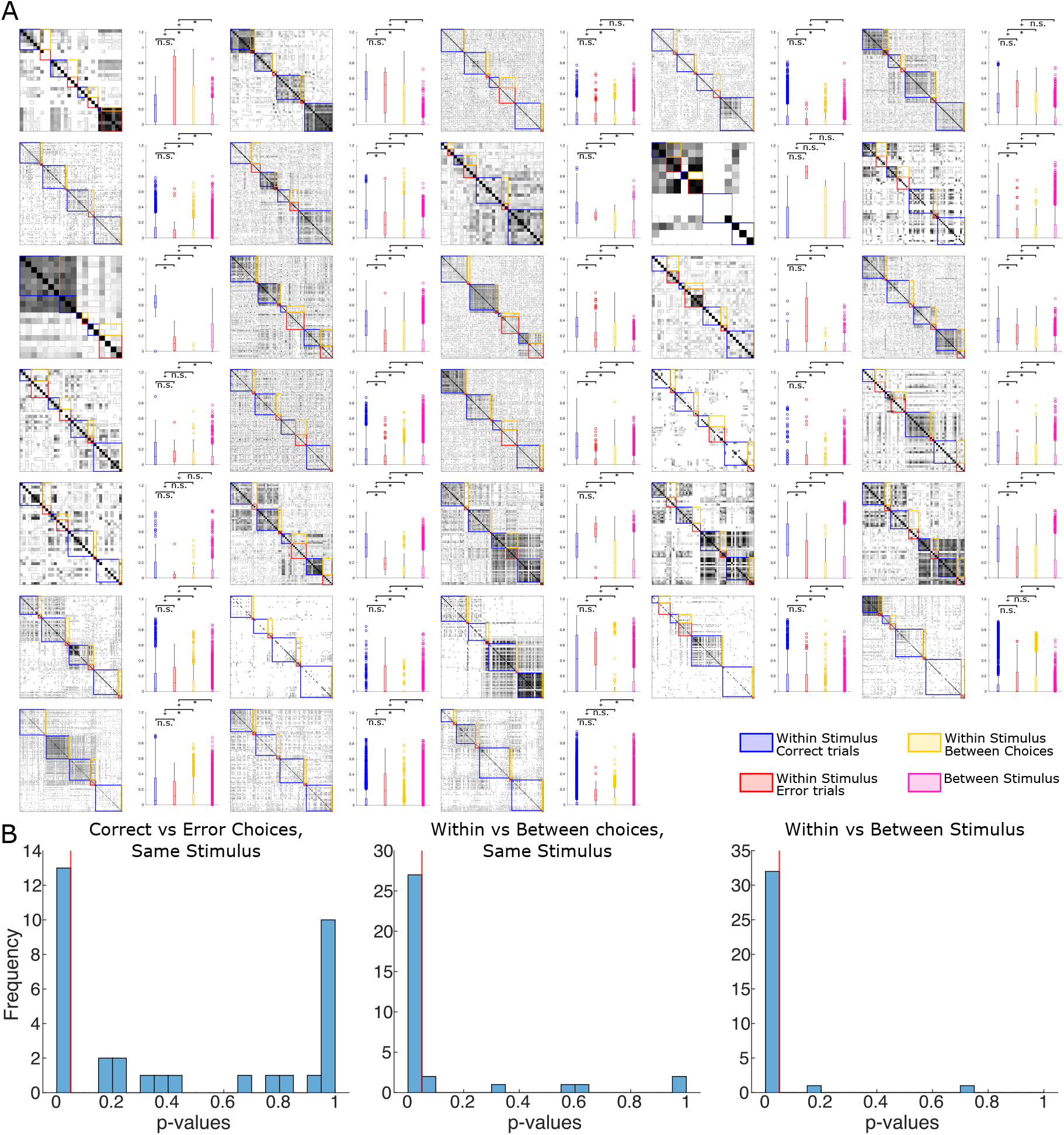
Trials Similarities in all sessions (A) Same similarity and test plots as in Fig 1B for each session. (B) Distribution of the p values of the three different tests conducted (correct vs error trials of same stimulus, within vs between choices of same stimulus, within vs between stimulus) in all the sessions.

**Figure S2.**
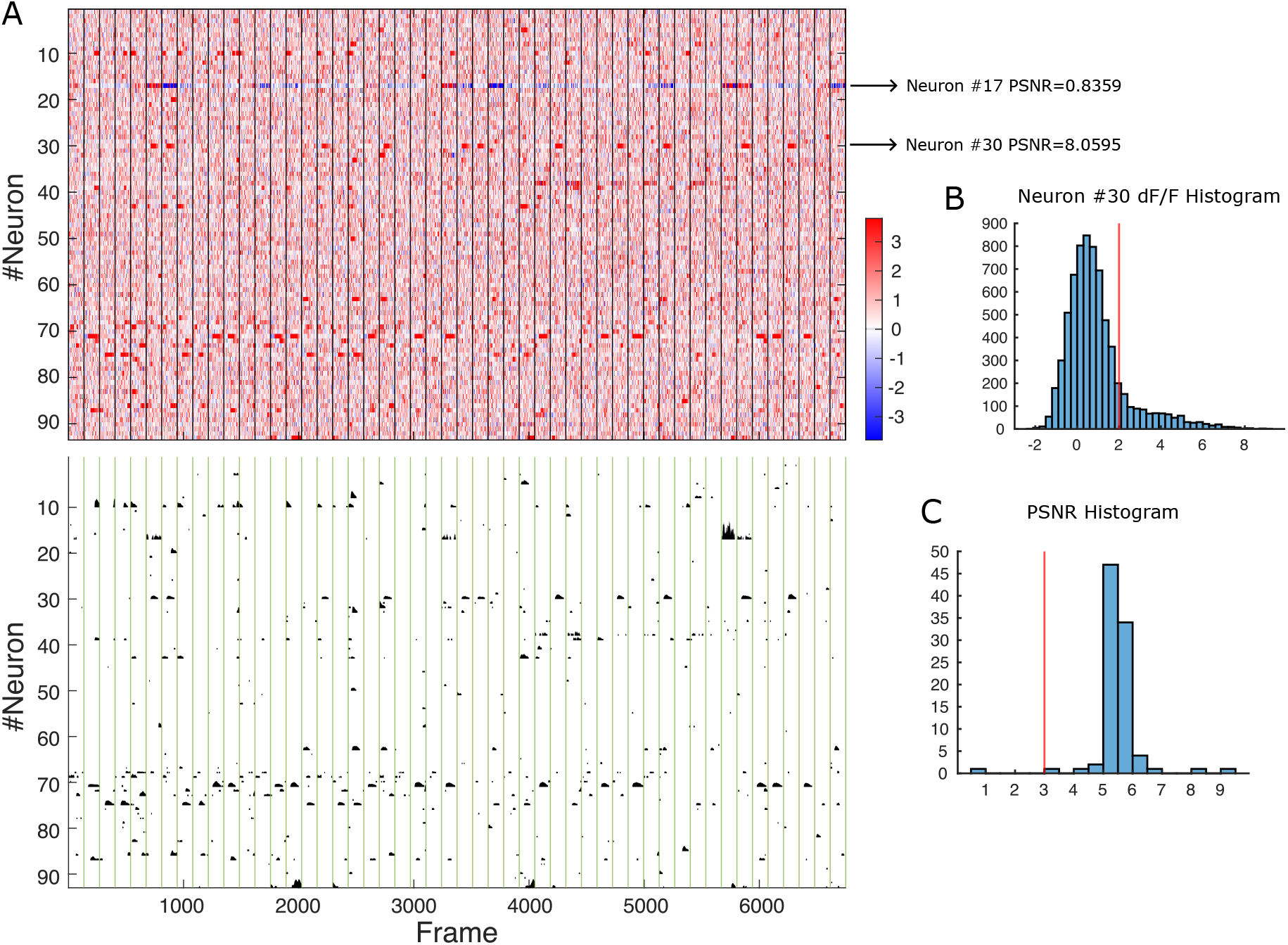
Preprocessing pipeline (A) Top: The dF/F traces of all the neurons in one example session. Each row is one neuron its activity was normalized to have unit robust standard deviation. Trials are segmented by vertical lines. PSNR was estimated for each neuron. The arrows indicate one highly responsive neuron (#30) and one corrupted neuron (#17). Bottom: keep only the true calcium transients in the session. (B) Distribution of dF/F of one responsive neuron (#30). A threshold of 2 (red line) was set to detect true transients. (C) Distribution of the PSNR of all the neurons. A threshold of 3 (red line) was set to filter out bad neurons.

**Figure S3A.**
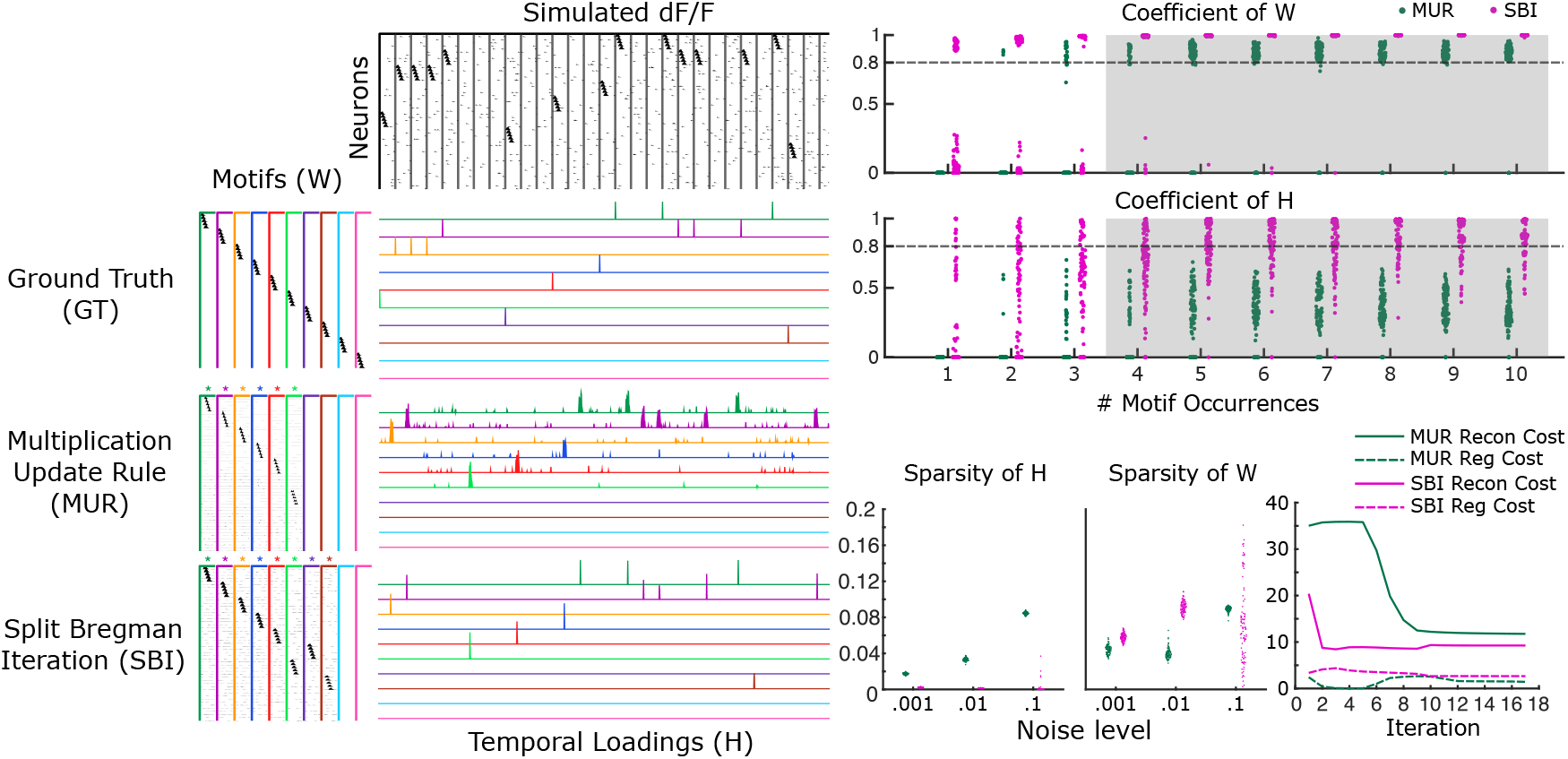
Compare SBI with MUR on simulated data (additive noise) Same plots as Fig 2B,C, running under additive noise condition(neurons are activated randomly beyond motif time courses). The activation probability of each neuron is set to 0.01 in the coefficient plots.

**Figure S3B.**
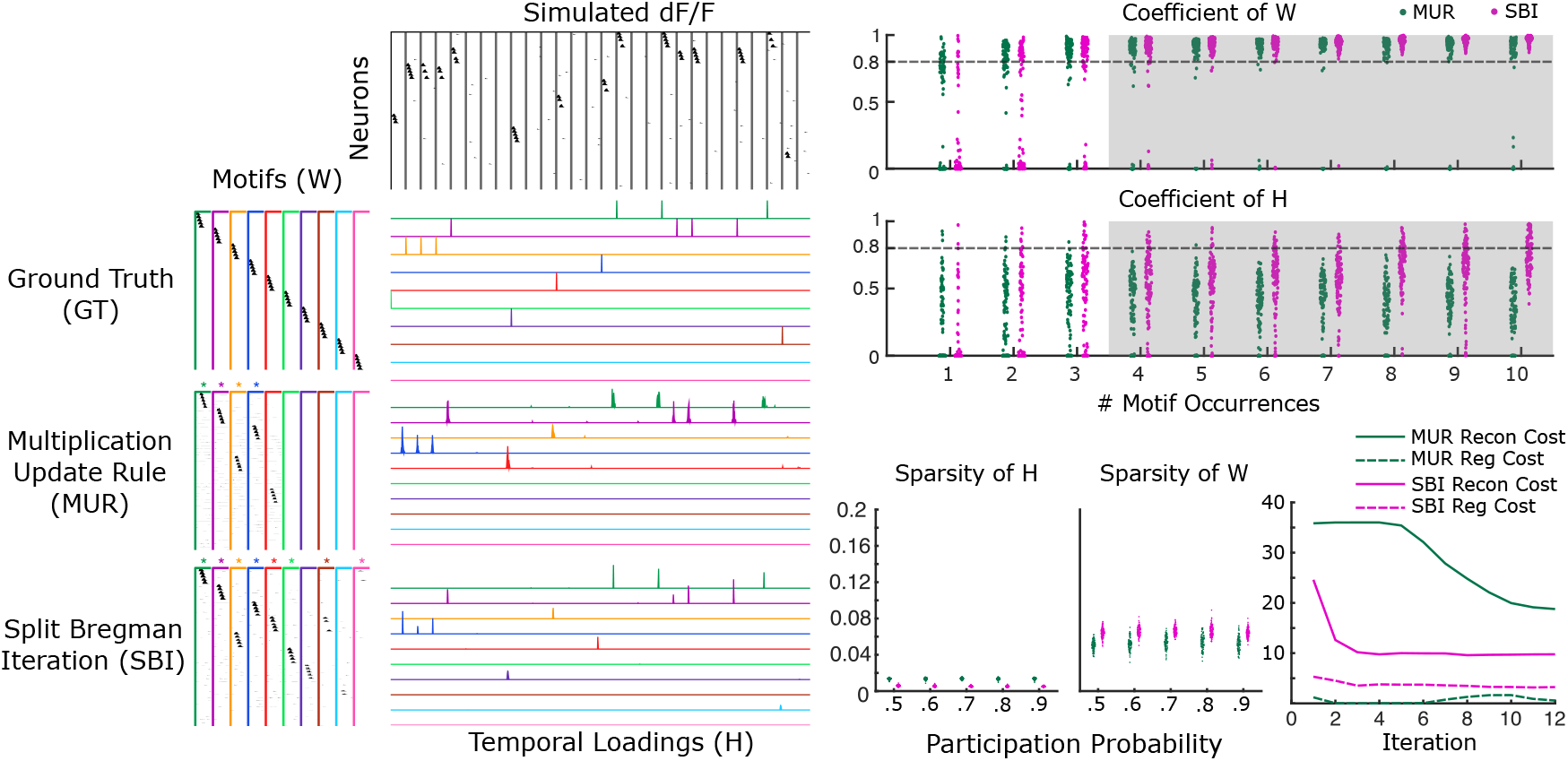
Compare SBI with MUR on simulated data (probabilistic participation) Same plots as Fig 2B,C, running under probabilistic participation condition(individual neurons participate with probability in each instance of the motif). The participation probability of each neuron is set to 0.7 in the coefficient plots.

**Figure S3C.**
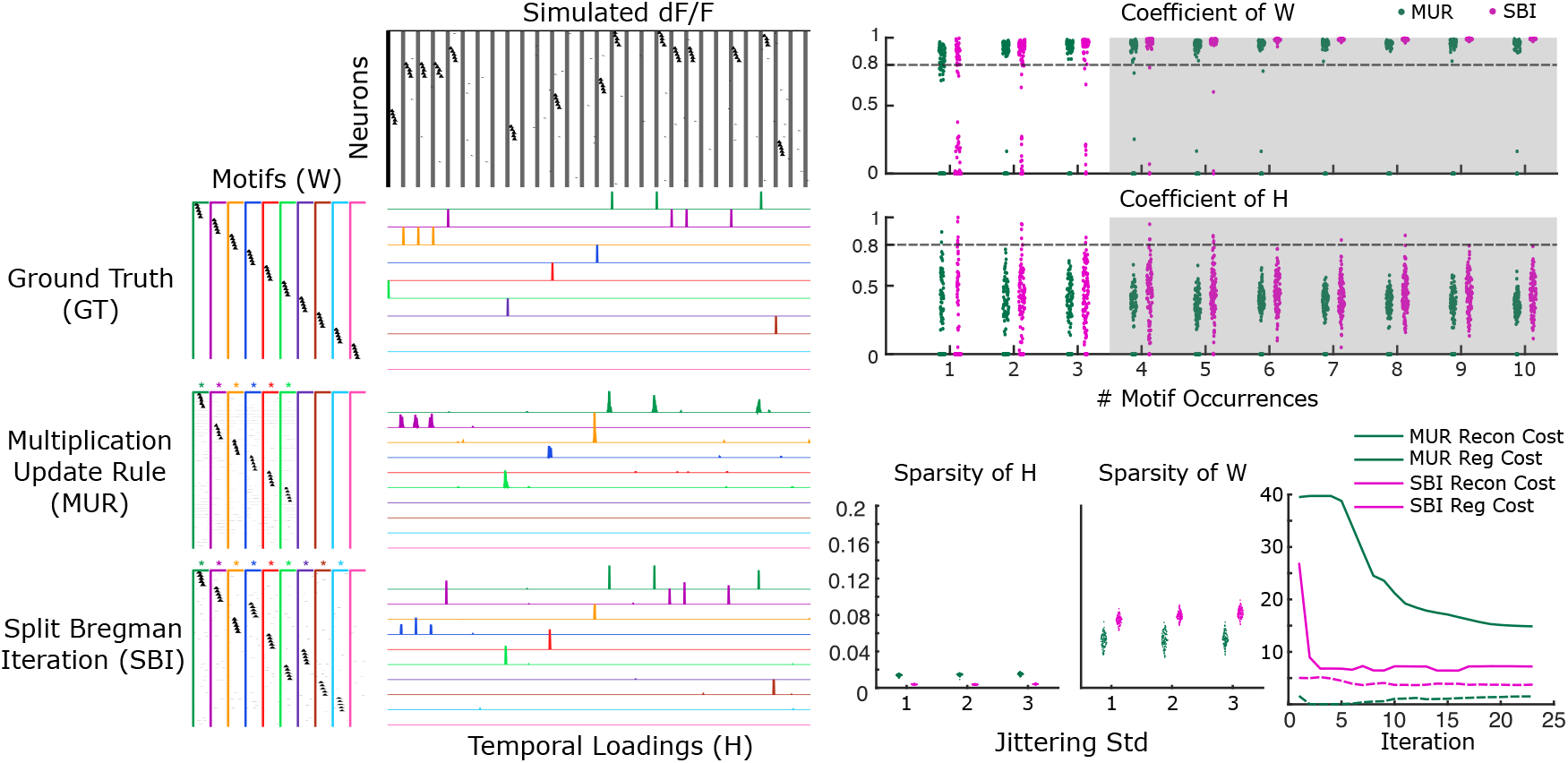
Compare SBI with MUR on simulated data (temporal jittering) Same plots as Fig 2B,C, running under temporal jittering condition (The activation time of each neuron is randomly shifted in each occurrence of the motif). The standard deviation of jittering time of each motif is set to 2 in the coefficient plots.

**Figure S3D.**
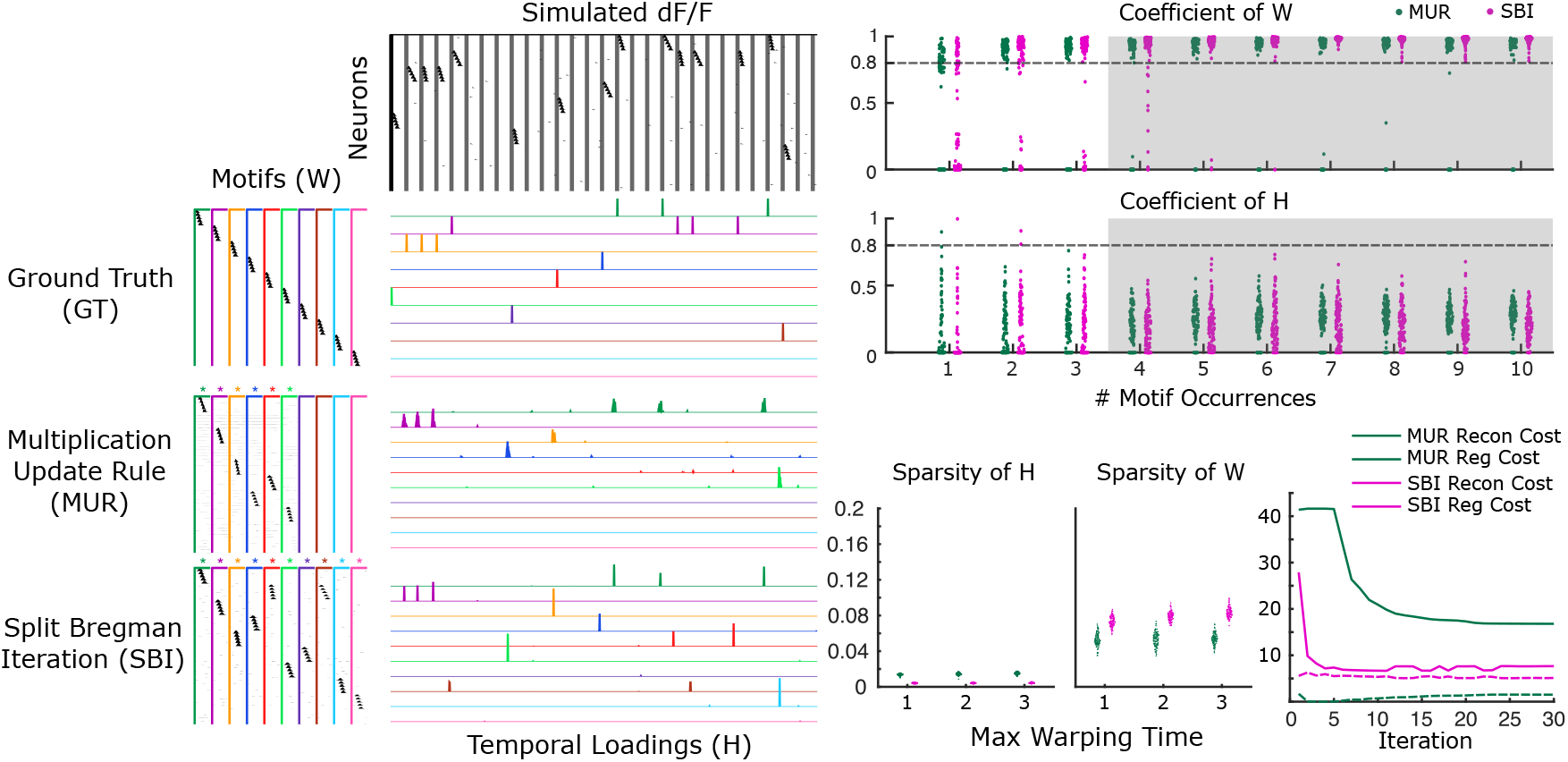
Compare SBI with MUR on simulated data (temporal warping) Same plots as Fig 2B,C, running under temporal warping condition (each instance of the motif occurs at a random speed). The max warping time unit of each motif is set to 2 in the coefficient plots.

**Figure S4.**
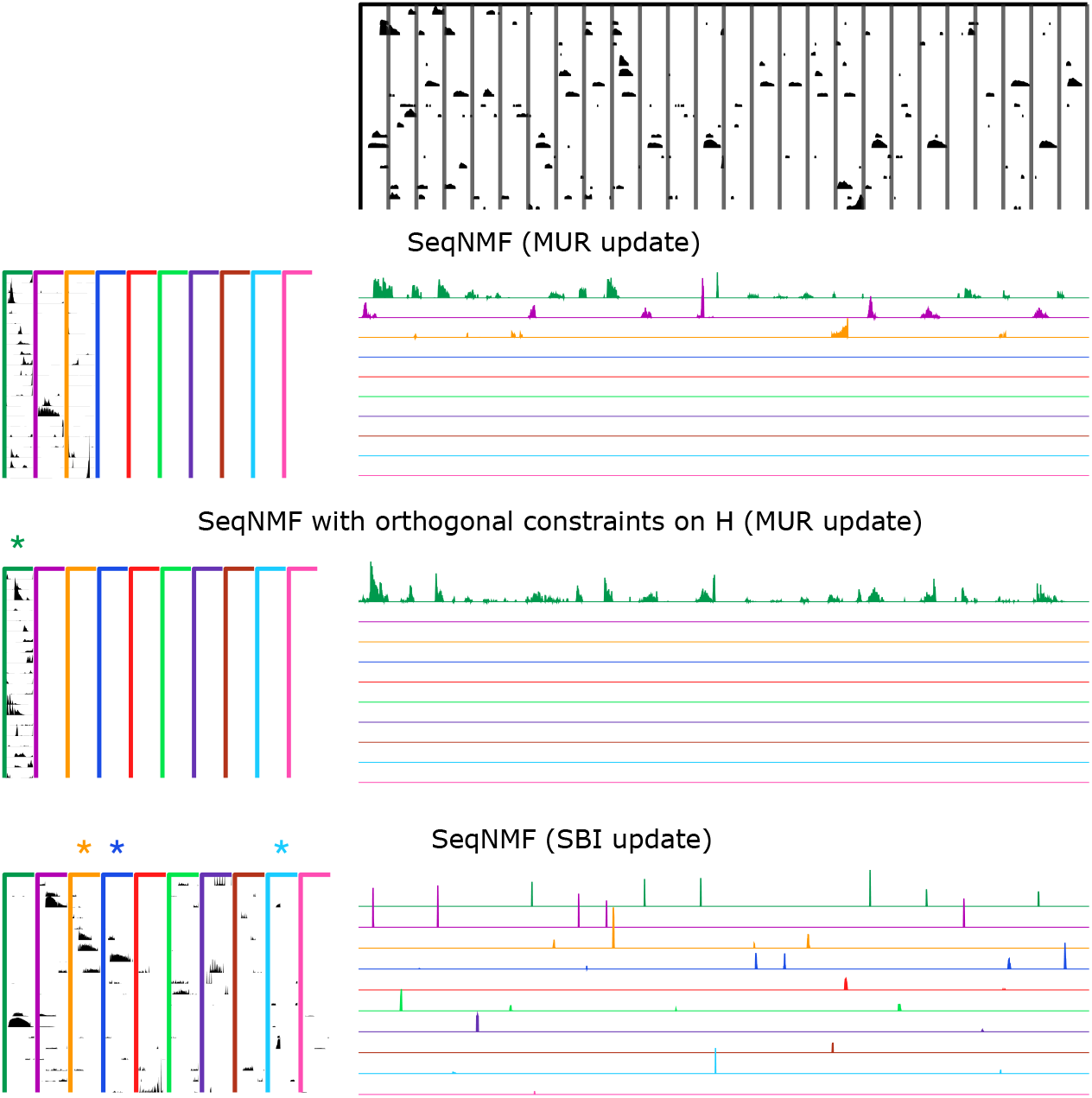
Adding Orthogonal regularization of H in SeqNMF does not improve temporal precision as using MUR Top: trainging data. Each row is the motifs and their temporal loading identified by different methods. To produce precise timing of **H**, we tried adding an additional orthogonal term of **H** in the SeqNMF optimization program (*λ* = 0.001, *λ*_*OrthH*_ = 0.01). Running with MUR method, SeqNMF could not produce the same precise temporal precision of **H** as using SBI. In addition, SBI was able to extract more motifs than MUR.

**Figure S5.**
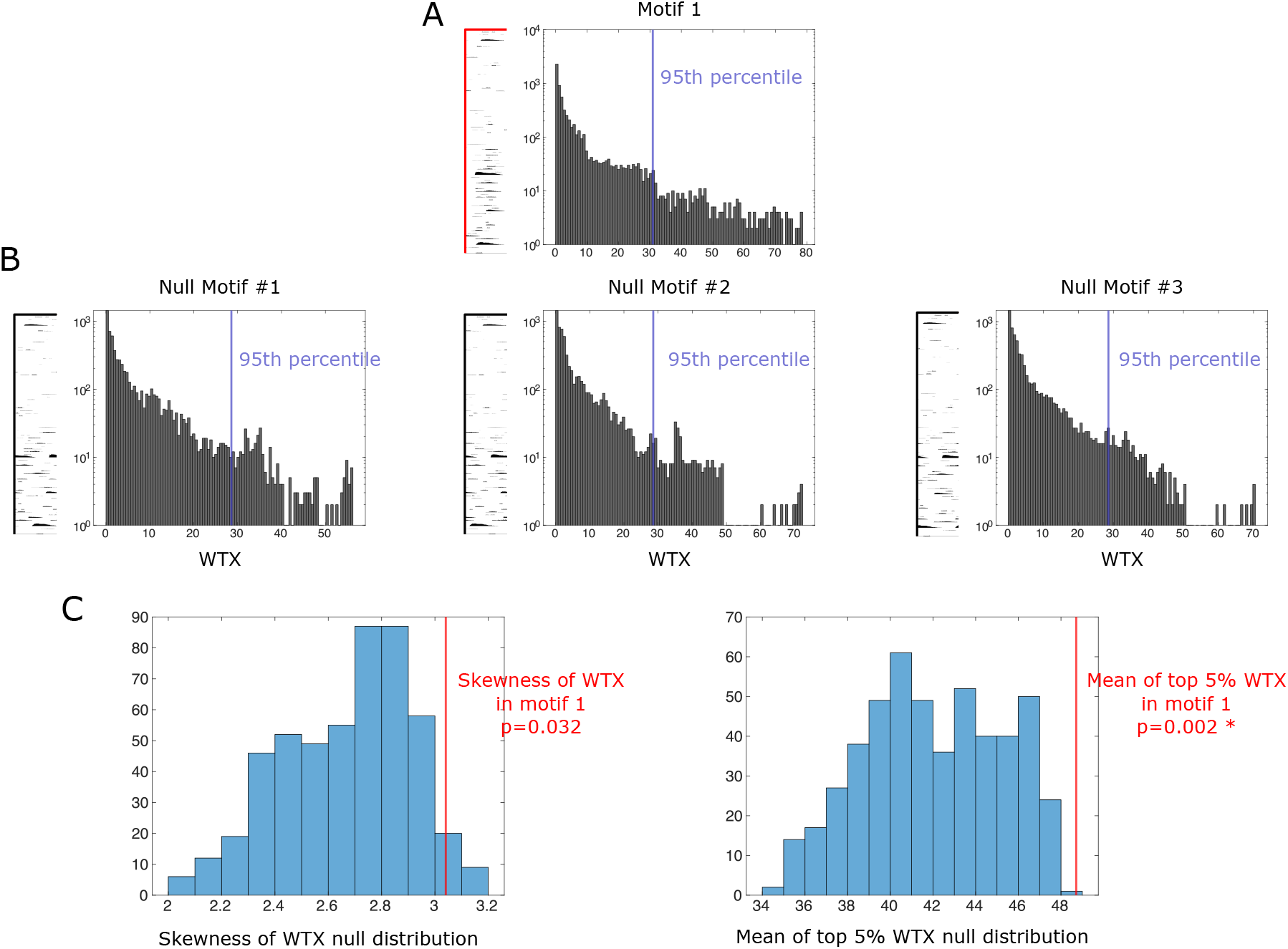
Significance test of motifs Adapted significance test of motifs based on SeqNMF paper. (A) Distribution of the overlap value 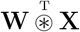 (denoted as WTX in the figure) between a motif and test data. (B) Null motifs were constructed by randomly circularly shifting each row of a motif independently. Multiple null motifs were constructed and the distribution of overlap values 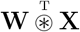 was measured between each null motif and the test data. The mean of the top 5% of the overlap values are calculated for each null motif and the original motif. (C) The mean of the top 5% of the overlap values is a more sensitive statistic to test the significance of the motif than the skewness of the overlap proposed by the SeqNMF paper. A motif is deemed significant if its mean of the top 5% of overlap values are greater than than of the null motifs.

**Figure S6.**
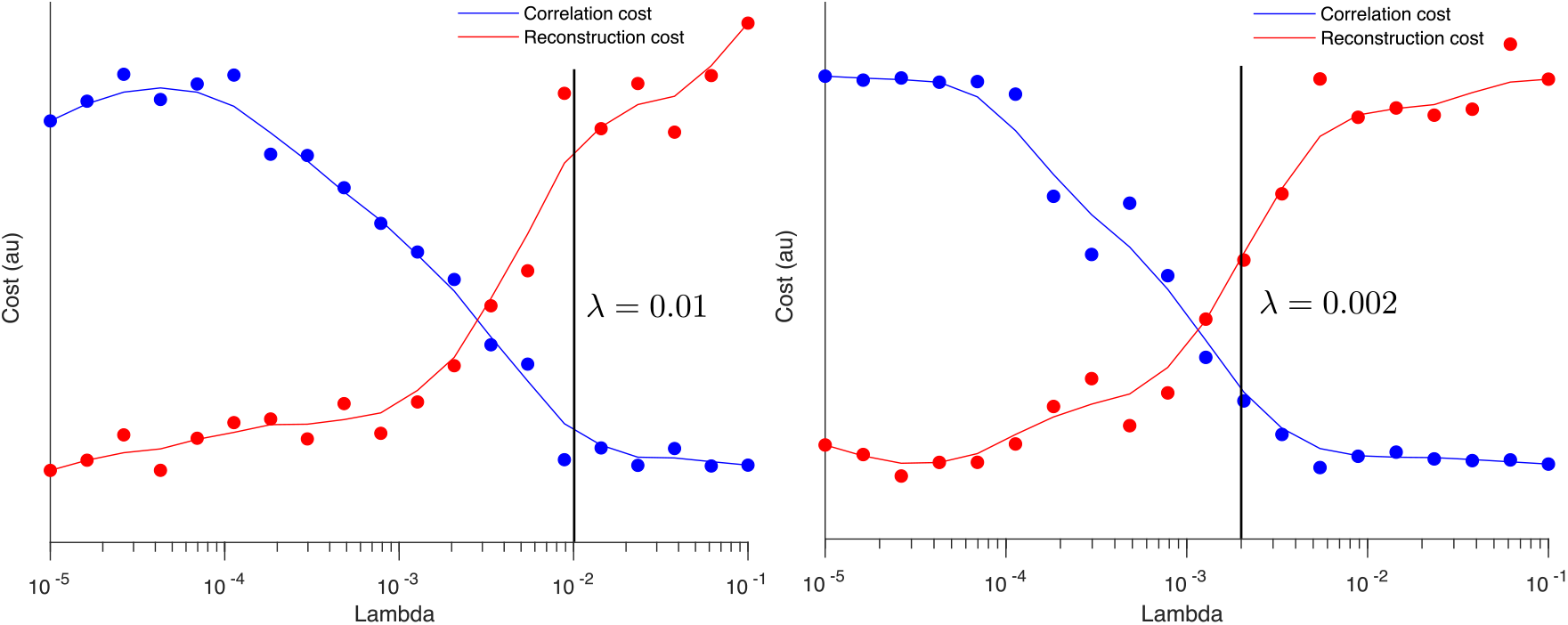
Choose optimal *λ* in SeqNMF Selection of optimal hyperparameter *λ* in SeqNMF (running with MUR). Left: Simulated data (neuron overlap condition). Right: Real data. Optimal *λ* was chosen (indicated by the black line) to be a few times larger than the crossing-point between the correlation cost and reconstruction cost.

**Figure S7.**
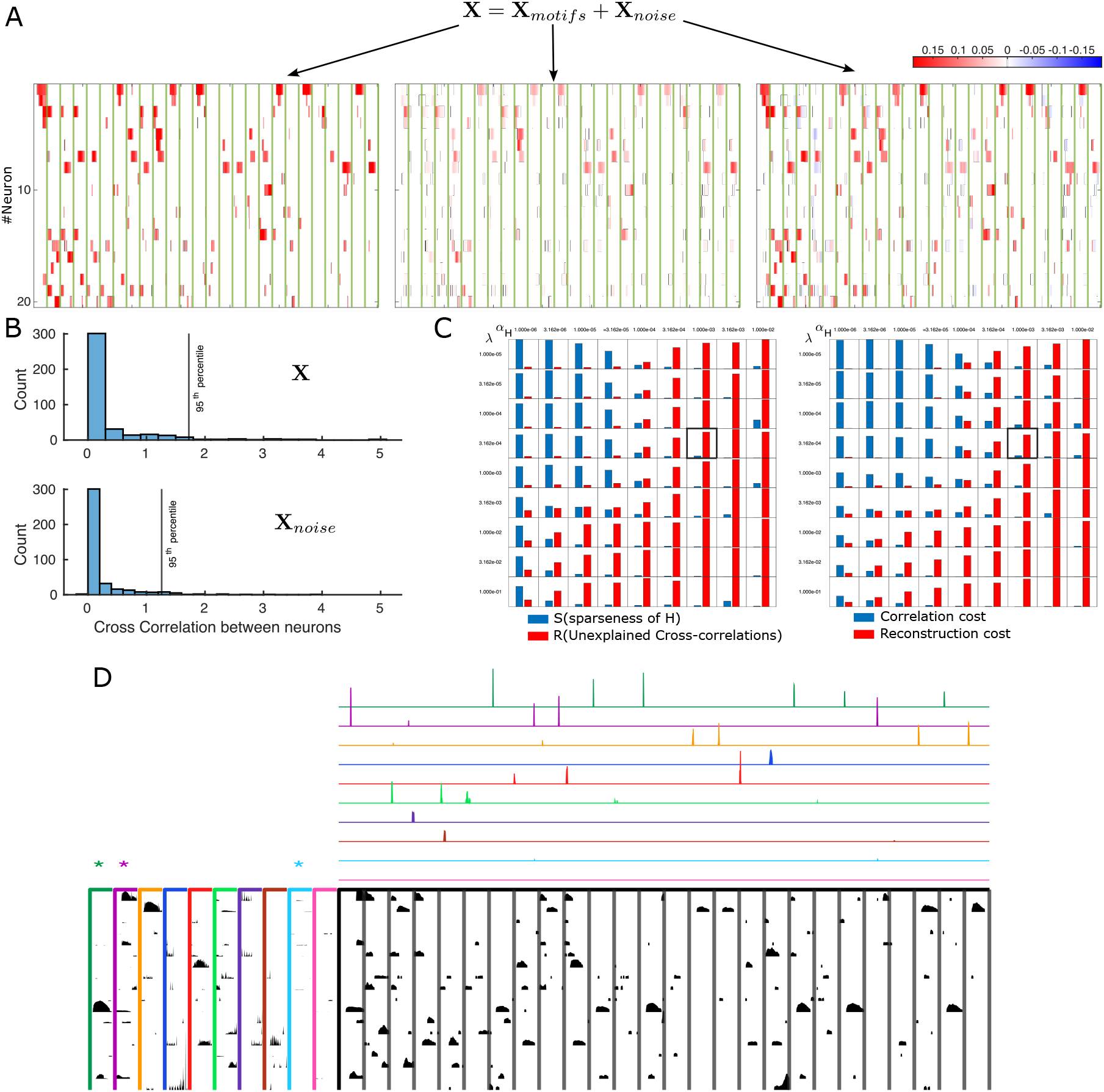
Selection of optimal hyperparameters in SBI (A) Decomposition of data **X** into stereotypical spatial-temporal motifs **X**_*motifs*_ and noise **X**_*noise*_. (B) If the motif structures are captured by the algorithm, the distribution of the cross-correlations between pairs of neurons in **X**_*noise*_ should move leftwards compared to **X**. (C) Plot of S(L0-norm of **H**) and R(the proportion of ‘motif’ structure in **X**_*noise*_) as a function of different hyperparameters *α*_**H**_ and *λ*. Bold bounding box indicates the best hyperparameters selected, which satisfy the optimal condition (Reconstruction cost slightly larger than regularization cost) in the SeqNMF procedure. (D) Running SBI to find motifs under the optimal hyperparameters selected above. Significant motifs are marked with asterisks.

**Figure S8.**
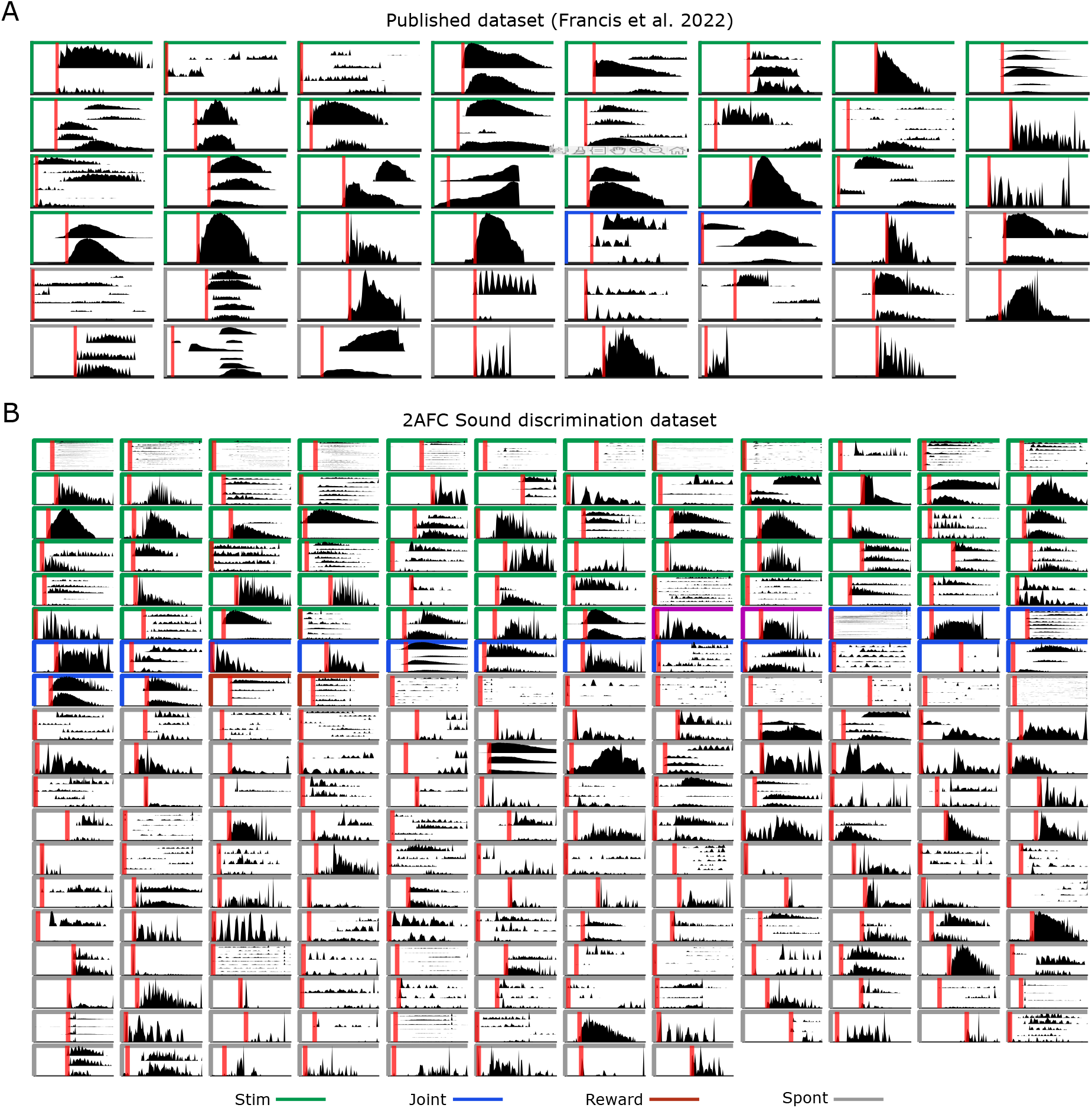
Task-encoding motif visualization (A) Visualization of all the task-encoding motifs and spontaneous motifs in Francis et al. dataset. (B) Same as (A) but on our own dataset.

**Figure S9.**
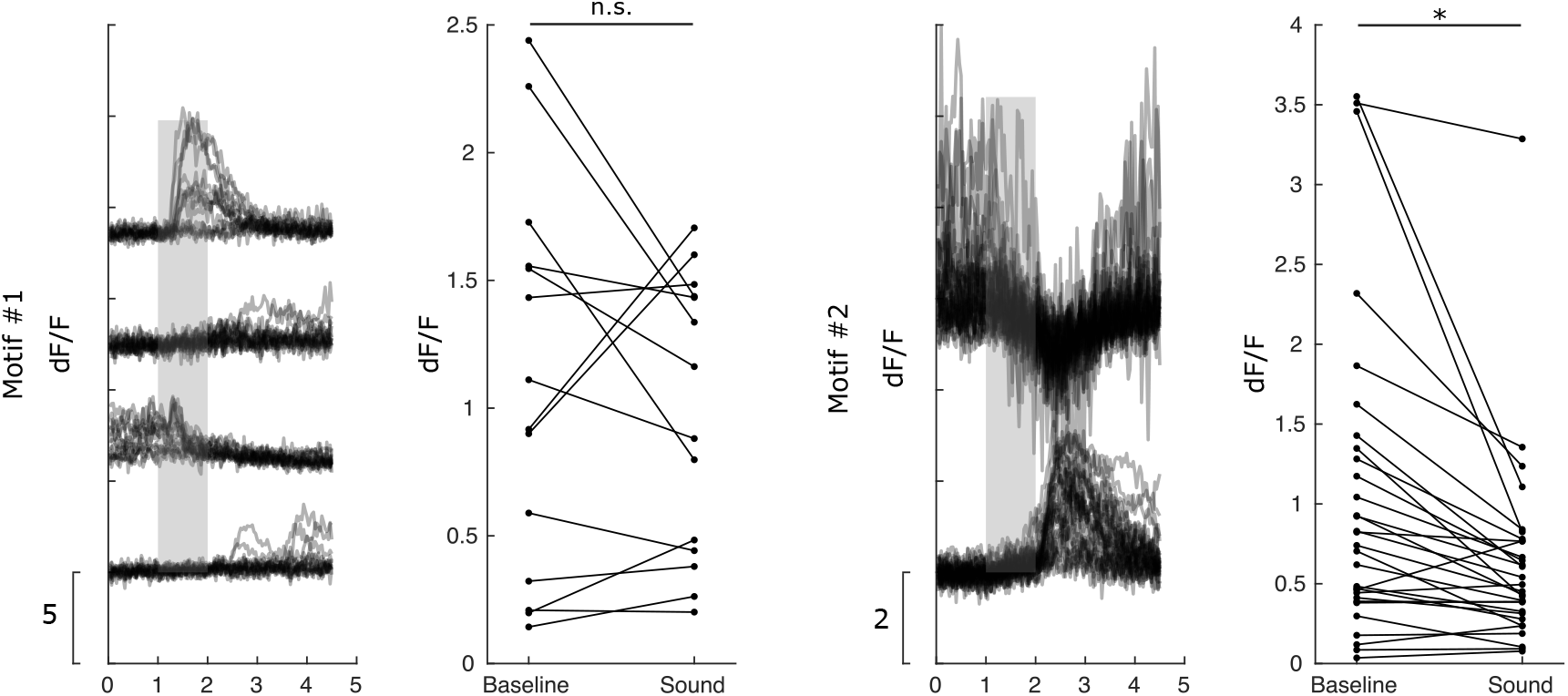
Early onsets of task-encoding motifs are caused by spontaneous activities before sound. Two example sound-encoding motifs with pre-sound onsets. Left panels show the dF/F traces of motif’s active neurons in all the trials with the encoding stimulus, right panels show a comparison between the mean dF/F over baseline and mean dF/F over sound in all the trials (paired t-test). The spontaneous activities of early-onset neurons (the third neuron in Motif #1 and the first neuron in Motif #2) can be greater or comparable to their activities during sound. However, they both exhibit greater spontaneous activities which are inhibited after sound in some trials, making them recruited by the motifs. Motif #1: p=0.41, Motif #2: p=0.001, paired two-sample ttest.

**Figure S10.**
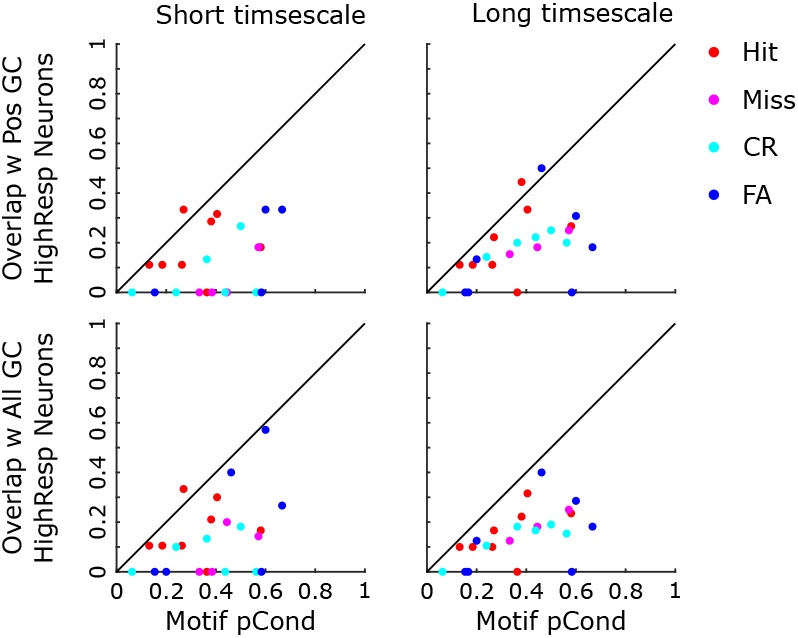
Comparing motif neurons with high-responsive GC-linked neurons Comparing motif neurons against GC-linked most responsive neurons (similar to Fig 4F).

**Figure S11.**
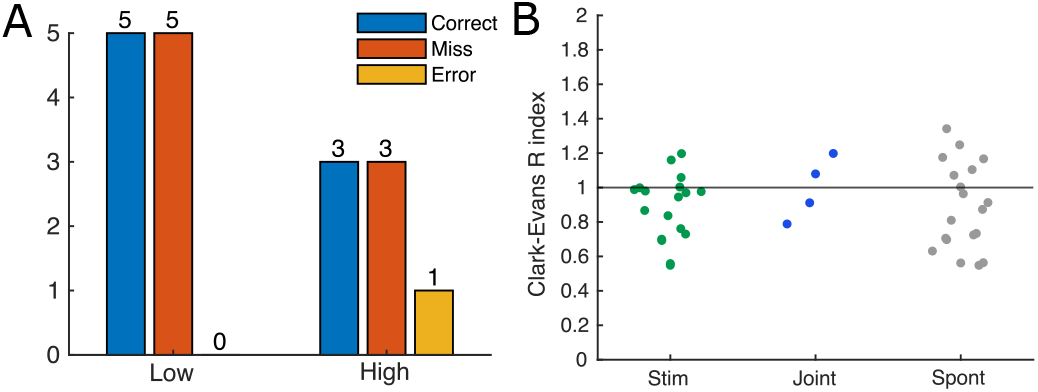
Properties of joint encoding motifs in 2AFC sound discrimination task (A) Encoding types of all the joint encoding motifs. Most motifs encode correct or error choice with a stimulus jointly. (B) Distribution of the Clark-Evan R index for all the stimulus-encoding, joint-encoding and spontaneous motifs. None of these indices significantly deviates from 1, indicating spatially uniformly distributed motif neurons.

**Figure S12.**
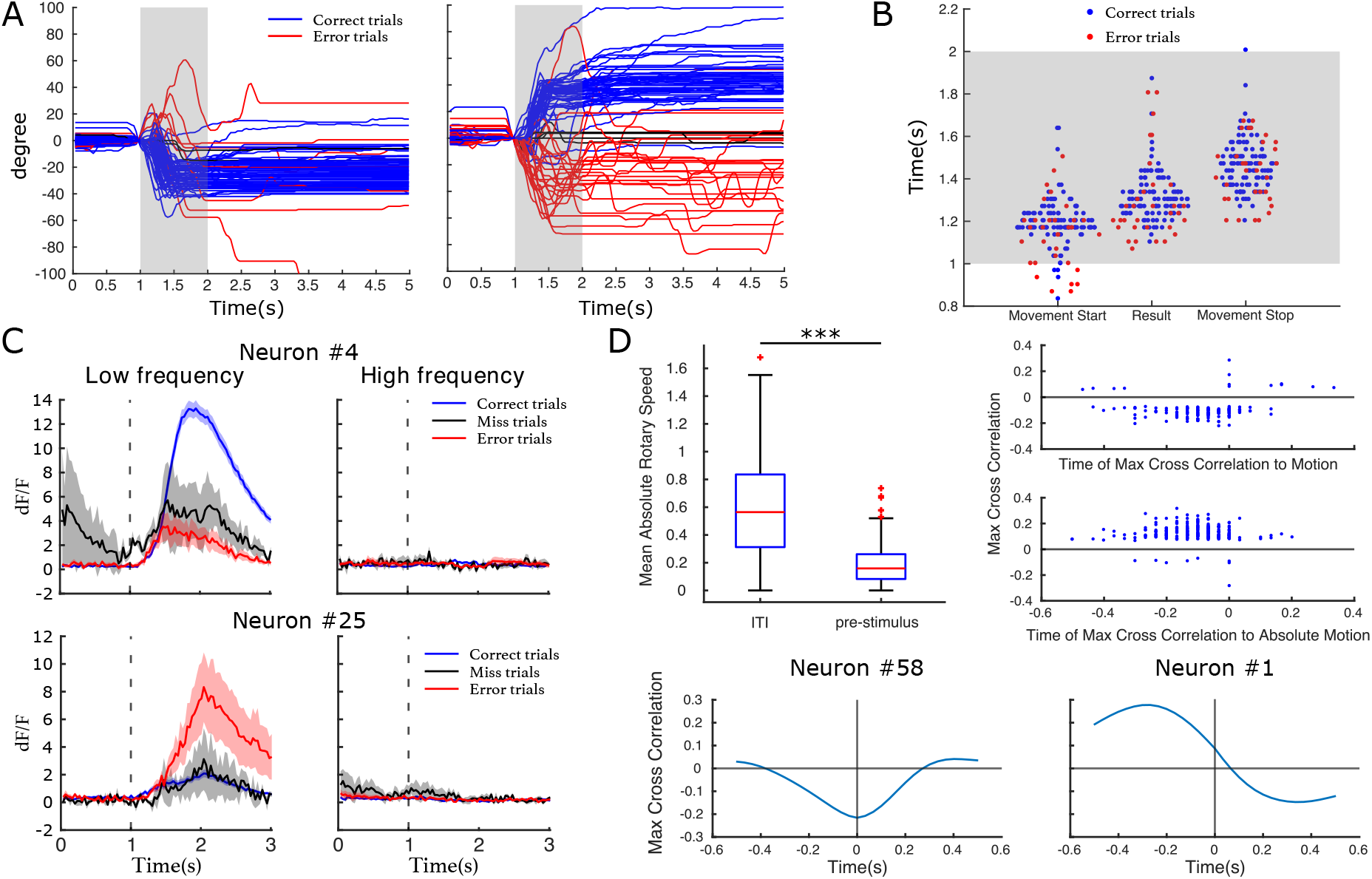
Single neuron encoding of task variables (A) Wheel movement trajectories in all the low (left) and high (right)frequency trials of one session. Blue: correct trials; red: error trials (B) Distribution of the time of movement onsets, trial results and movement offsets in the session in A. Note that in some trials there are spontaneous movement before trial onsets, leading to movement start or stop events before sound. (C) Two example neurons jointly encoding sound and choice. For each neuron we plotted the mean (solid line) and s.e.m (shaded region) of the neuron’s dF/F activities in different trials. Top: one neuron encoding low frequency and correct trials jointly. Bottom: one neuron encoding low frequency and error trials jointly. Vertical dashed lines: sound onsets. (D) Top Left: Significantly greater wheel movement activities were observed during the inter-trial intervals (ITI) compared to 1 second before sound onsets. We thus used the neural activities during inter-trial intervals to test the encoding of wheel movement. Top right: distribution of max cross correlations and their peak times. Bottom left: Cross correlations between movement and neuron activity at different time lags for one example neuron. Bottom right: Cross correlations between absolute movement and neuron activity at different time lags for another example neuron.

**Figure S13.**
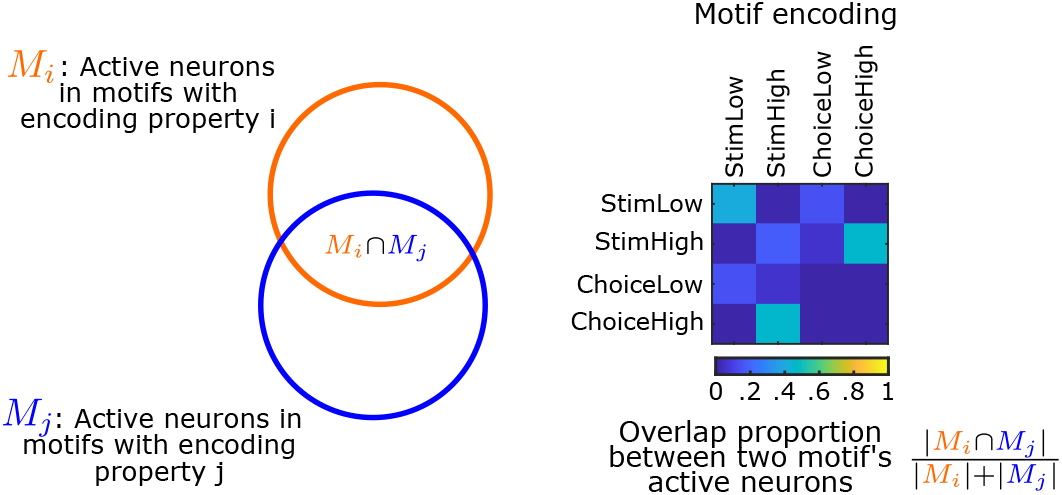
Overlap between active neurons of different motifs Mean of the overlaps (measure as dice coefficients) between active neurons of two different motifs, grouped by motifs’ encoding.

## References

[1] Eberhard Zwicker and Karin Schorn. Psychoacoustical tuning curves in audiology. Audiology, 17(2):120–140, 1978.

[2] NYS Kiang and EC Moxon. Tails of tuning curves of auditory-nerve fibers. The Journal of the Acoustical Society of America, 55(3):620–630, 1974.

[3] Daniel A Butts and Mark S Goldman. Tuning curves, neuronal variability, and sensory coding. PLoS biology, 4(4):e92, 2006.

[4] Apostolos P Georgopoulos and Costas N Stefanis. Local shaping of function in the motor cortex: motor contrast, directional tuning. Brain research reviews, 55(2):383–389, 2007.

[5] Andrew D Huberman, Marla B Feller, and Barbara Chapman. Mechanisms underlying development of visual maps and receptive fields. Annu. Rev. Neurosci., 31(1):479–509, 2008.

[6] Moshe Abeles. Transmission of Information by Coincidence. In Moshe Abeles, editor, Local Cortical Circuits: An Electrophysiological Study, Studies of Brain Function, pages 67–75. Springer, Berlin, Heidelberg, 1982.

[7] Yuji Ikegaya, Gloster Aaron, Rosa Cossart, Dmitriy Aronov, Ilan Lampl, David Ferster, and Rafael Yuste. Synfire Chains and Cortical Songs: Temporal Modules of Cortical Activity. Science, 304(5670):559–564, April 2004.

[8] Artur Luczak, Bruce L. McNaughton, and Kenneth D. Harris. Packet-based communication in the cortex. Nature Reviews Neuroscience, 16(12):745–755, December 2015.

[9] Luis Carrillo-Reid, Weijian Yang, Jae-eun Kang Miller, Darcy S. Peterka, and Rafael Yuste. Imaging and Optically Manipulating Neuronal Ensembles. Annual Review of Biophysics, 46(Volume 46, 2017):271–293, May 2017.

[10] Rafael Yuste, Rosa Cossart, and Emre Yaksi. Neuronal ensembles: Building blocks of neural circuits. Neuron, 112(6):875–892, March 2024.

[11] Artur Luczak, Peter Barthó, and Kenneth D. Harris. Spontaneous Events Outline the Realm of Possible Sensory Responses in Neocortical Populations. Neuron, 62(3):413–425, May 2009.

[12] Luis Carrillo-Reid, Shuting Han, Weijian Yang, Alejandro Akrouh, and Rafael Yuste. Controlling Visually Guided Behavior by Holographic Recalling of Cortical Ensembles. Cell, 178(2):447–457.e5, July 2019.

[13] Richard H. R. Hahnloser, Alexay A. Kozhevnikov, and Michale S. Fee. An ultra-sparse code underliesthe generation of neural sequences in a songbird. Nature, 419(6902):65–70, September 2002.

[14] Tatsuo S Okubo, Emily L Mackevicius, Hannah L Payne, Galen F Lynch, and Michale S Fee. Growth and splitting of neural sequences in songbird vocal development. Nature, 528(7582):352–357, 2015.

[15] Zoltán Nádasdy, Hajime Hirase, András Czurkó, Jozsef Csicsvari, and György Buzsáki. Replay and Time Compression of Recurring Spike Sequences in the Hippocampus. Journal of Neuroscience, 19(21):9497–9507, November 1999.

[16] Eva Pastalkova, Vladimir Itskov, Asohan Amarasingham, and György Buzsáki. Internally Generated Cell Assembly Sequences in the Rat Hippocampus. Science, 321(5894):1322–1327, September 2008.

[17] Thomas J. Davidson, Fabian Kloosterman, and Matthew A. Wilson. Hippocampal Replay of Extended Experience. Neuron, 63(4):497–507, August 2009.

[18] Christopher D. Harvey, Philip Coen, and David W. Tank. Choice-specific sequences in parietal cortex during a virtual-navigation decision task. Nature, 484(7392):62–68, April 2012.

[19] Lloyd E. Russell, Mehmet Fişek, Zidan Yang, Lynn Pei Tan, Adam M. Packer, Henry W. P. Dalgleish, Selmaan N. Chettih, Christopher D. Harvey, and Michael Häusser. The influence of cortical activity on perception depends on behavioral state and sensory context. Nature Communications, 15(1):2456, March 2024.

[20] Raphael Steinfeld, André Tacão-Monteiro, and Alfonso Renart. Differential representation of sensory information and behavioral choice across layers of the mouse auditory cortex. Current Biology, 34(10):2200–2211.e6, May 2024.

[21] David M Schneider. Reflections of action in sensory cortex. Current Opinion in Neurobiology, 64:53–59, October 2020.

[22] Matthias Minderer, Kristen D. Brown, and Christopher D. Harvey. The Spatial Structure of Neural Encoding in Mouse Posterior Cortex during Navigation. Neuron, 102(1):232–248.e11, April 2019.

[23] Kishore V. Kuchibhotla, Jonathan V. Gill, Grace W. Lindsay, Eleni S. Papadoyannis, Rachel E. Field, Tom A. Hindmarsh Sten, Kenneth D. Miller, and Robert C. Froemke. Parallel processing by cortical inhibition enables context-dependent behavior. Nature Neuroscience, 20(1):62–71, January 2017.

[24] Mu Zhou, Feixue Liang, Xiaorui R. Xiong, Lu Li, Haifu Li, Zhongju Xiao, Huizhong W. Tao, and Li I. Zhang. Scaling down of balanced excitation and inhibition by active behavioral states in auditory cortex. Nature Neuroscience, 17(6):841–850, June 2014.

[25] Ningyu Zhang and Ning-long Xu. Reshaping sensory representations by task-specific brain states: Toward cortical circuit mechanisms. Current Opinion in Neurobiology, 77:102628, December 2022.

[26] Nicholas A. Steinmetz, Peter Zatka-Haas, Matteo Carandini, and Kenneth D. Harris. Distributed coding of choice, action and engagement across the mouse brain. Nature, 576(7786):266–273, December 2019.

[27] Shinichiro Kira, Houman Safaai, Ari S. Morcos, Stefano Panzeri, and Christopher D. Harvey. A distributed and efficient population code of mixed selectivity neurons for flexible navigation decisions. Nature Communications, 14(1):2121, April 2023.

[28] Camden J. MacDowell and Timothy J. Buschman. Low-Dimensional Spatiotemporal Dynamics Underlie Cortex-wide Neural Activity. Current Biology, 30(14):2665–2680.e8, July 2020.

[29] Nikolas A Francis, Shoutik Mukherjee, Loren Koçillari, Stefano Panzeri, Behtash Babadi, and Patrick O Kanold. Sequential transmission of task-relevant information in cortical neuronal networks. Cell reports, 39(9):110878, May 2022.

[30] Nikolas A. Francis, Daniel E. Winkowski, Alireza Sheikhattar, Kevin Armengol, Behtash Babadi, and Patrick O. Kanold. Small Networks Encode Decision-Making in Primary Auditory Cortex. Neuron, 97(4):885–897.e6, February 2018.

[31] Sue Ann Koay, Adam S. Charles, Carlos D. Brody, and David W. Tank. Sequential and efficient neural-population coding of complex task information. Neuron, 110(2):328–349.e11, January 2022.

[32] Paris Smaragdis. Non-negative Matrix Factor Deconvolution; Extraction of Multiple Sound Sources from Monophonic Inputs. In Carlos G. Puntonet and Alberto Prieto, editors, Independent Component Analysis and Blind Signal Separation, Lecture Notes in Computer Science, pages 494–499, Berlin, Heidelberg, 2004. Springer.

[33] Zhe Chen and Andrzej Cichocki. Nonnegative matrix factorization with temporal smoothness and/or spatial decorrelation constraints. Signal Processing, November 2004.

[34] Paul D. O’Grady and Barak A. Pearlmutter. Convolutive Non-Negative Matrix Factorisation with a Sparseness Constraint. In 2006 16th IEEE Signal Processing Society Workshop on Machine Learning for Signal Processing, pages 427–432, September 2006.

[35] Vikram Ramanarayanan, Louis Goldstein, and Shrikanth S. Narayanan. Spatio-temporal articulatory movement primitives during speech production: Extraction, interpretation, and validation. The Journal of the Acoustical Society of America, 134(2):1378–1394, August 2013.

[36] Alex H. Williams, Anthony Degleris, Yixin Wang, and Scott W. Linderman. Point process models for sequence detection in high-dimensional neural spike trains. Advances in neural information processing systems, 33:14350–14361, December 2020.

[37] Emily L Mackevicius, Andrew H Bahle, Alex H Williams, Shijie Gu, Natalia I Denisenko, Mark S Goldman, and Michale S Fee. Unsupervised discovery of temporal sequences in high-dimensional datasets, with applications to neuroscience. eLife, 8:e38471, February 2019.

[38] Michael W. Berry, Murray Browne, Amy N. Langville, V. Paul Pauca, and Robert J. Plemmons. Algorithms and applications for approximate nonnegative matrix factorization. Computational Statistics & Data Analysis, 52(1):155–173, September 2007.

[39] Yinan Li, Ruili Wang, Yuqiang Fang, Meng Sun, and Zhangkai Luo. Alternating Direction Method of Multipliers for Convolutive Non-Negative Matrix Factorization. IEEE Transactions on Cybernetics, pages 1–14, 2022.

[40] Dennis L. Sun and Cédric Févotte. Alternating direction method of multipliers for non-negative matrix factorization with the beta-divergence. In 2014 IEEE International Conference on Acoustics, Speech and Signal Processing (ICASSP), pages 6201–6205, May 2014.

[41] Rafal Zdunek and Andrzej Cichocki. Nonnegative matrix factorization with constrained second-order optimization. Signal Processing, 87(8):1904–1916, August 2007.

[42] Andrzej Cichocki, Rafal Zdunek, and Shun-ichi Amari. Csiszár’s Divergences for Non-negative Matrix Factorization: Family of New Algorithms. In Justinian Rosca, Deniz Erdogmus, José C. Príncipe, and Simon Haykin, editors, Independent Component Analysis and Blind Signal Separation, Lecture Notes in Computer Science, pages 32–39, Berlin, Heidelberg, 2006. Springer.

[43] Anh Huy Phan, Andrzej Cichocki, Petr Tichavský, and Zbyněk Koldovský. On Connection between the Convolutive and Ordinary Nonnegative Matrix Factorizations. In Fabian Theis, Andrzej Cichocki, Arie Yeredor, and Michael Zibulevsky, editors, Latent Variable Analysis and Signal Separation, Lecture Notes in Computer Science, pages 288–296, Berlin, Heidelberg, 2012. Springer.

[44] Jian-Feng Cai, Stanley Osher, and Zuowei Shen. Split Bregman Methods and Frame Based Image Restoration. Multiscale Modeling & Simulation, 8(2):337–369, January 2010.

[45] Tom Goldstein and Stanley Osher. The Split Bregman Method for L1-Regularized Problems. SIAM Journal on Imaging Sciences, 2(2):323–343, January 2009.

[46] Daniel Lee and H. Sebastian Seung. Algorithms for Non-negative Matrix Factorization. In Advances in Neural Information Processing Systems, volume 13. MIT Press, 2000.

[47] Daniel D. Lee and H. Sebastian Seung. Learning the parts of objects by non-negative matrix factorization. Nature, 401(6755):788–791, October 1999.

[48] Philip J. Clark and Francis C. Evans. Distance to Nearest Neighbor as a Measure of Spatial Relationships in Populations. Ecology, 35(4):445–453, 1954.

[49] Shreya Saxena and John P Cunningham. Towards the neural population doctrine. Current Opinion in Neurobiology, 55:103–111, April 2019.

[50] Stefano Panzeri, Christopher D. Harvey, Eugenio Piasini, Peter E. Latham, and Tommaso Fellin. Cracking the Neural Code for Sensory Perception by Combining Statistics, Intervention, and Behavior. Neuron, 93(3):491–507, February 2017.

[51] Giuseppe Pica, Eugenio Piasini, Houman Safaai, Caroline Runyan, Christopher Harvey, Mathew Diamond, Christoph Kayser, Tommaso Fellin, and Stefano Panzeri. Quantifying how much sensory information in a neural code is relevant for behavior. In Advances in Neural Information Processing Systems, volume 30. Curran Associates, Inc., 2017.

[52] C. W. J. Granger. Investigating Causal Relations by Econometric Models and Cross-spectral Methods. Econometrica, 37(3):424–438, 1969.

[53] Mingxuan Wang, Peter Jendrichovsky, and Patrick O. Kanold. Auditory discrimination learning differentially modulates neural representation in auditory cortex subregions and inter-areal connectivity. Cell Reports, 43(5), May 2024.

[54] Hiroaki Tsukano, Masao Horie, Takeshi Bo, Arikuni Uchimura, Ryuichi Hishida, Masaharu Kudoh, Kuniyuki Takahashi, Hirohide Takebayashi, and Katsuei Shibuki. Delineation of a frequency-organized region isolated from the mouse primary auditory cortex. Journal of Neurophysiology, 113(7):2900–2920, April 2015.

[55] Jonathan W. Pillow, Jonathon Shlens, Liam Paninski, Alexander Sher, Alan M. Litke, E. J. Chichilnisky, and Eero P. Simoncelli. Spatio-temporal correlations and visual signalling in a complete neuronal population. Nature, 454(7207):995–999, August 2008.

[56] Ji Liu, Matthew R Whiteway, Alireza Sheikhattar, Daniel A Butts, Behtash Babadi, and Patrick O Kanold. Parallel processing of sound dynamics across mouse auditory cortex via spatially patterned thalamic inputs and distinct areal intracortical circuits. Cell reports, 27(3):872– 885, 2019.

[57] Ziyue Aiden Wang, Susu Chen, Yi Liu, Dave Liu, Karel Svoboda, Nuo Li, and Shaul Druckmann. Not everything, not everywhere, not all at once: A study of brain-wide encoding of movement, June 2023.

[58] Anton Filipchuk, Joanna Schwenkgrub, Alain Destexhe, and Brice Bathellier. Awake perception is associated with dedicated neuronal assemblies in the cerebral cortex. Nature Neuroscience, 25(10):1327–1338, October 2022.

[59] Daniel E Winkowski, Daniel A Nagode, Kevin J Donaldson, Pingbo Yin, Shihab A Shamma, Jonathan B Fritz, and Patrick O Kanold. Orbitofrontal Cortex Neurons Respond to Sound and Activate Primary Auditory Cortex Neurons. Cerebral Cortex, 28(3):868–879, March 2018.

[60] Jonah K. Mittelstadt and Patrick O. Kanold. Orbitofrontal cortex conveys stimulus and task information to the auditory cortex. Current Biology, 33(19):4160–4173.e4, October 2023.

[61] Lin Zhong, Yuan Zhang, Chunyu A. Duan, Ji Deng, Jingwei Pan, and Ning-long Xu. Causal contributions of parietal cortex to perceptual decision-making during stimulus categorization. Nature Neuroscience, 22(6):963–973, June 2019.

[62] Anders Nelson, David M. Schneider, Jun Takatoh, Katsuyasu Sakurai, Fan Wang, and Richard Mooney. A Circuit for Motor Cortical Modulation of Auditory Cortical Activity. Journal of Neuroscience, 33(36):14342–14353, September 2013.

[63] Ye Liang, Jing Li, Yu Tian, Peng Tang, Chunhua Liu, and Xi Chen. The Anterior Cingulate Cortex Promotes Long-Term Auditory Cortical Responses through an Indirect Pathway via the Rhinal Cortex in Mice. Journal of Neuroscience, 43(23):4262–4278, June 2023.

[64] Kenneth D. Harris and Alexander Thiele. Cortical state and attention. Nature Reviews Neuroscience, 12(9):509–523, September 2011.

[65] Ji Liu and Patrick O. Kanold. Interactive auditory task reveals complex sensory-action integration in mouse primary auditory cortex, December 2022.

[66] Theda Backen, Stefan Treue, and Julio C. Martinez-Trujillo. Encoding of Spatial Attention by Primate Prefrontal Cortex Neuronal Ensembles. eNeuro, 5(1), January 2018. Publisher: Society for Neuroscience Section: New Research.

[67] Jermyn Z See, Craig A Atencio, Vikaas S Sohal, and Christoph E Schreiner. Coordinated neuronal ensembles in primary auditory cortical columns. eLife, 7:e35587, June 2018.

[68] Congcong Hu, Andrea R. Hasenstaub, and Christoph E. Schreiner. Basic Properties of Coordinated Neuronal Ensembles in the Auditory Thalamus. Journal of Neuroscience, 44(19), May 2024.

[69] Mohamad Abbass, Benjamin Corrigan, Renée Johnston, Roberto Gulli, Adam Sachs, Jonathan C. Lau, and Julio Martinez-Trujillo. Neural Ensembles in the Lateral Prefrontal Cortex Temporally Multiplex Task Features During Virtual Navigation, January 2024.

[70] Shumpei P. Yasuda, Yuta Seki, Sari Suzuki, Yasuhiro Ohshiba, Xuehan Hou, Kunie Matsuoka, Kenta Wada, Hiroshi Shitara, Yuki Miyasaka, and Yoshiaki Kikkawa. C.753A>G genome editing of a Cdh23 ahl allele delays age-related hearing loss and degeneration of cochlear hair cells in C57BL/6J mice. Hearing Research, 389:107926, April 2020.

[71] Ji Liu and Patrick O. Kanold. Diversity of Receptive Fields and Sideband Inhibition with Complex Thalamocortical and Intracortical Origin in L2/3 of Mouse Primary Auditory Cortex. Journal of Neuroscience, 41(14):3142–3162, April 2021.

[72] Tsai-Wen Chen, Trevor J. Wardill, Yi Sun, Stefan R. Pulver, Sabine L. Renninger, Amy Baohan, Eric R. Schreiter, Rex A. Kerr, Michael B. Orger, Vivek Jayaraman, Loren L. Looger, Karel Svoboda, and Douglas S. Kim. Ultrasensitive fluorescent proteins for imaging neuronal activity. Nature, 499(7458):295–300, July 2013.

[73] Jeffrey L. Gauthier, Sue Ann Koay, Edward H. Nieh, David W. Tank, Jonathan W. Pillow, and Adam S. Charles. Detecting and correcting false transients in calcium imaging. Nature Methods, 19(4):470–478, April 2022.

[74] Peter J. Huber. Robust Statistics. John Wiley & Sons, 2004.

[75] Stephen R. Becker, Emmanuel J. Candés, and Michael C. Grant. Templates for convex cone problems with applications to sparse signal recovery. Mathematical Programming Computation, 3(3):165, July 2011.

[76] Alfred Auslender and Marc Teboulle. Interior Gradient and Proximal Methods for Convex and Conic Optimization. SIAM Journal on Optimization, 16(3):697–725, January 2006.

[77] Y. Nesterov. A method for unconstrained convex minimization problem with the rate of convergence o(1/k2). 1983.

[78] Yu. Nesterov. Smooth minimization of non-smooth functions. Mathematical Programming, 103(1):127–152, May 2005.

[79] Amir Beck and Marc Teboulle. A Fast Iterative Shrinkage-Thresholding Algorithm for Linear Inverse Problems. SIAM Journal on Imaging Sciences, 2(1):183–202, January 2009.

[80] Yu. Nesterov. Gradient methods for minimizing composite functions. Mathematical Programming, 140(1):125–161, August 2013.

